# Spatial protein and RNA analysis on the same tissue section using MICS technology

**DOI:** 10.1101/2023.10.27.564191

**Authors:** Emily Neil, Dongju Park, Rebecca C. Hennessey, Eric C. DiBiasio, Michael DiBuono, Hanna Lafayette, Erica Lloyd, Hsinyi Lo, Julia Femel, Alex Makrigiorgos, Sameh Soliman, Dominic Mangiardi, Paurush Praveen, Silvia Rüberg, Fabian Staubach, Ryan Hindman, Thomas Rothmann, Hansueli Meyer, Tanya Wantenaar, Jinling Wang, Werner Müller, Robert Pinard, Andreas Bosio

## Abstract

Spatial Biology has evolved from the molecular characterization of microdissected cells to high throughput spatial RNA and protein expression analysis at scale. The main limitation of spatial technologies so far is the inability to resolve protein and RNA information in the same histological section. Here, we report for the first time the integration of highly multiplexed RNA and protein detection on the same tissue section. We developed a new, automated, spatial RNA detection method (RNAsky™), which is based on targeted rolling circle amplification and iterative staining. We combine RNAsky with MACSima™ Imaging Cyclic Staining (MICS) based protein analysis and show compatibility with subsequent standard hematoxylin and eosin (H&E) staining. Using both, open-source tools and our recently developed software suite MACS® iQ View, we demonstrate our multiomics MICS workflow by characterizing key immune-oncology markers at subcellular resolution across normal and diseased tissues.

## Introduction

Understanding the composition and architecture of cells within tissues is fundamental to studying tissue function, and more broadly, human biology. The complex dynamics of cellular functions and interactions between cell populations drive normal physiology, with dysregulation of these mechanisms underlying disease progression. Advances in bulk and single cell sequencing have facilitated the establishment of cell atlas initiatives, which aim to map all cells in the human body in healthy and diseased states ^1, 2, 3, 4, 5, 6, 7, 8, 9,10^. Although sequencing methods allow for transcriptome wide analyses of individual cells with high sensitivity and accuracy, they require dissociation of cells, thereby destroying all contextual information ^11, 12^.

Image-based, spatially resolved transcriptomics methods have been extensively reviewed, and fall primarily into two categories: fluorescent *in situ* hybridization (FISH) based or rolling circle amplification (RCA) based approaches ^13, 14, 15, 16, 17^. With FISH based approaches, complementary probes are hybridized to mRNAs of interest, and repeatedly interrogated to generate optical barcodes associated with each gene ^18, 19, 20, 21, 22, 23^. While optical barcoding allows for high multiplexing, it requires long imaging times and advanced algorithms for barcode decoding and downstream analyses. For RCA approaches, a circular template is required for amplification and is achieved either by using targeted padlock probes or circularizing templates *in situ* ^24, 25, 26, 27, 28, 29^. Rolling circle amplicons can be interrogated in a number of ways, including sequencing by ligation or hybridization-based readouts. A combination of low plex *in situ* fluorescence mRNA analysis with MALDI mass spectrometry imaging or immunofluorescent protein detection has been reported in literature ^30, 31, 32^. However, spatially resolved and highly multiplexed RNA detection methods have so far shown limited compatibility for same-slide iterative immunofluorescence-based protein detection methods.

Spatial proteomics is a vital counterpart to spatial transcriptomics, as it provides direct assessments of functional information. Numerous multiplex proteomics technologies have been developed, including iterative and simultaneous detection methods ^33, 34, 35, 36, 37, 38^. One such advance is MACSima™ imaging cyclic staining (MICS), which is a fully automated, cyclic immunofluorescence technology performed on the MACSima platform utilizing directly conjugated antibodies combined with gentle signal erasure methods, thereby allowing the application of hundreds of antibodies to a single sample ^33^. Due to the non-destructive nature of this process, the MACSima platform is an ideal system for combining multiplex proteomics with spatial transcriptomics.

Here, we introduce RNAsky™, a targeted, RCA-based gene expression profiling assay designed for the MACSima platform. We combined this assay with multiplex protein imaging to generate multiomic datasets in normal and diseased tissues. To demonstrate the capabilities of our multiomics platform, we profiled multiple tissues using a targeted RNAsky Immuno-Oncology (I-O) Core panel with additional customized gene targets and a selection of protein markers.

## Results

### The MACSima Platform enables same-section multiplex analysis of proteins and mRNA

A comprehensive understanding of the complex interactions between cells in a spatial context requires multiple information modalities. Recently, MICS technology was introduced for spatial analysis of epitopes such as proteins, glycolipids, and complex sugars at subcellular resolution using hundreds of immunofluorescent antibodies on the same specimen ^33^. To add the capability of transcript detection, we developed RNAsky, which is based on the same principle of iterative staining, sample washing, multi-field imaging, and signal erasure. Like protein detection, the RNAsky assay is performed on the MACSima Platform, a fully automated, fluorescence microscopy instrument. We specifically optimized transcript amplification and detection chemistries to be compatible with subsequent epitope detection in order to combine the analysis of RNA and protein targets on a single formalin-fixed paraffin-embeded (FFPE) tissue section (Fig. 1a, Methods). We chose a workflow starting with RNA target probe amplification, followed by iterative detection probe hybridization, and ending with immunofluorescent detection of epitopes. Similar to directly-conjugated antibodies, each gene displayed a unique emission spectrum during a single cycle via the hybridization of gene-specific detection probes. Transcripts were detected across four wavelengths (FITC, PE, APC, near infrared), and proteins were detected with three wavelengths (FITC, PE, APC). This technology supports simultaneous analyses of protein markers and RNA transcripts, enabling the characterization of the spatial architecture and expression topography of tissues.

**Fig. 1.**
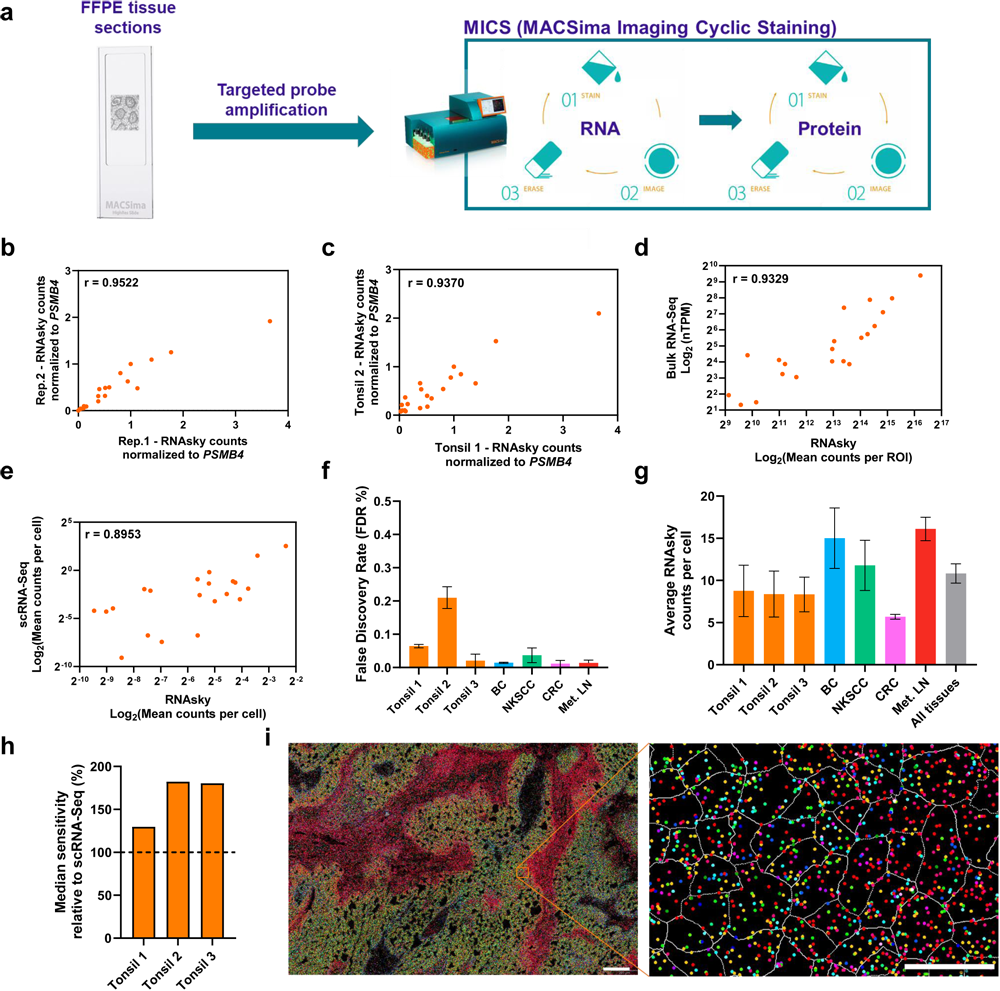
Schematic workflow and performance of RNAsky. **a** Overview of MACSima Imaging Cyclic Staining (MICS), an automated, multiomic approach to integrate spatial protein and RNA analysis on a single tissue section. **b** Representative correlation of RNAsky counts between independent runs (serial section replicates) of a representative normal tonsil tissue. Rep. 1 vs Rep. 2 Pearson correlation coefficient (PCC) = 0.9522 (95% confidence intervals = 0.8865 to 0.9803); P value (two-tailed) <0.0001. Remaining correlations in Supplementary Fig. 1. **c** Correlation of RNAsky counts obtained in two different normal tonsil blocks (n=3 for each block). PCC = 0.9370 (95% confidence intervals = 0.8486 to 0.9745); P value (two-tailed) <0.0001. **d** Correlation of a normal tonsillar bulk RNA-Seq dataset (Human Protein Atlas) and RNAsky counts of a representative normal tonsil. PCC = 0.9329 (95% confidence intervals = 0.8392 to 0.9728); P value (two-tailed) <0.0001. **e** Correlation of a normal young adult tonsillar scRNA-Seq dataset and RNAsky counts of a representative normal tonsil. PCC = 0.8953 (95% confidence intervals = 0.7558 to 0.9571); P value (two-tailed) <0.0001. **f** False discovery rate (FDR) of RNAsky in all tissue (BC – breast cancer, NKSCC – non-keratinizing squamous cell carcinoma, CRC – colorectal cancer, Met. LN – metastatic lymph node) analyzed. Data represents mean ± SEM (n=3 for tonsil, BC, NKSCC, Met. LN; n=2 for CRC). **g** Average transcripts per cell in all tissues analyzed. Data represents mean ± SEM (n=3 for tonsil, BC, NKSCC, met. LN; n=2 for CRC; n = 20 for all tissues). **h** Median *in situ* sensitivity of RNAsky in three normal tonsils relative to a normal young adult tonsillar scRNA-Seq dataset. The line at 100% indicates the ratio where the sensitivity of scRNA-Seq and RNAsky are equal. **i** Subcellular resolution of RNAsky transcript detection in a representative NKSCC ROI. Colored dots indicate the genes in extended I-O panel and grey lines indicate cell segmentation. Image (scale bar = 200 µm; left) and close-up region (scale bar = 20 µm; right).

### RNAsky is a reproducible, sensitive, specific, and spatially resolved gene expression assay

RNAsky is a spatial gene expression assay designed to complement the proteomic capabilities of the MACSima platform. Briefly, the assay is based on targeted padlock probe hybridization, ligation, and amplification. Padlock probes are oligonucleotides whose ends are complementary to adjacent target sequences ^24, 39^. During hybridization, the two binding portions of the padlock must perfectly hybridize to the target to bring the 5’ and 3’ ends into contact, allowing circularization of the probes by ligation. Rolling circle amplification then generates multiple copies of the circularized probes, producing DNA nanoballs. To assess the performance of RNAsky, normal tonsil samples were interrogated with a Human I-O Core Panel, composed of 21 target genes, two housekeeping genes, and negative controls. Regions of interest (ROI) containing a minimum of two follicles across three serial sections were evaluated to determine intra-sample and inter-sample reproducibility, as well as cross technology comparison (Fig. 1b-e). To correct for sample-to-sample variation, RNAsky counts were normalized to the higher expressed housekeeping gene in the panel, *PSMB4*. RNAsky showed high concordance across serial sections of the same block (r=0.9522 or higher) (Fig. 1b, Supplementary Fig. 1a) with an average of 22,782 cells profiled across the three sections. RNAsky was also highly concordant between two independent tissue samples with similar regions of interest assayed (Tonsil 1= 22,782 cells, Tonsil 2 = 21,313 cells, r=0.9370) (Fig. 1c). We then compared RNAsky data with two independent RNA quantification technologies, bulk RNA-Seq and single-cell RNA-Seq to benchmark the assay’s performance (Fig. 1d-e, Supplementary Fig. 1b). RNAsky showed high correlation with both methods (Bulk RNA-Seq correlation r= 0.9329, scRNA-Seq correlation r= 0.8953).

We developed RNAsky as a flexible assay, where the number and composition of targets can be customized according to study needs. To highlight this flexibility and expand cell type characterization, four additional gene targets were added to the I-O Core Panel (*BCL6*, *CD68*, *CD14*, *LAMP3*, referred to as extended I-O panel) in cancer samples of interest. Specificity was evaluated by calculating the false discovery rate (FDR) across multiple replicates per tissue type (Fig. 1f). RNAsky demonstrated highly specific performance, with all FDR’s falling below 0.3%. The number of counts per cell varied across tissue types, with an average of 10.8 counts per cell in all tissues (Fig. 1g, Supplementary Fig. 1c). To benchmark the sensitivity of RNAsky, the ratio of mean counts per cell for each gene was calculated in both RNAsky and scRNA-Seq tonsil data (Supplementary Fig. 1d). RNAsky showed higher sensitivity for low and medium expressed genes, but this sensitivity dropped for highly expressed transcripts. For highly expressed genes, switching from a discrete count-based quantification to continuous intensity-based quantification provided a robust alternative to assess transcript abundance (Supplementary Fig. 2). The median sensitivity was calculated for all genes across three tonsil blocks, and showed higher sensitivity compared to scRNA-Seq (Fig. 1h). Overall, RNAsky provides spatially resolved quantification of transcripts at subcellular resolution with high reproducibility, specificity, and sensitivity (Fig. 1i).

### The combination of RNA and protein analyses identifies major cell types within normal tonsil tissue

The ability to capture tissue architecture on a transcriptional level was investigated by assessing the gene expression patterns of the extended I-O panel in normal tonsil (Fig. 2). The spatial distribution of detected genes demonstrated major characteristics of tonsil histology, including the germinal center and marginal zone of lymphoid follicles (Fig. 2, Region 1), the interfollicular zone (Fig. 2, Region 2), and squamous epithelium (Fig. 2, Region 3). The patterns of gene expression were localized across all panel targets (Supplementary Figs. 3-5).

**Fig. 2.**
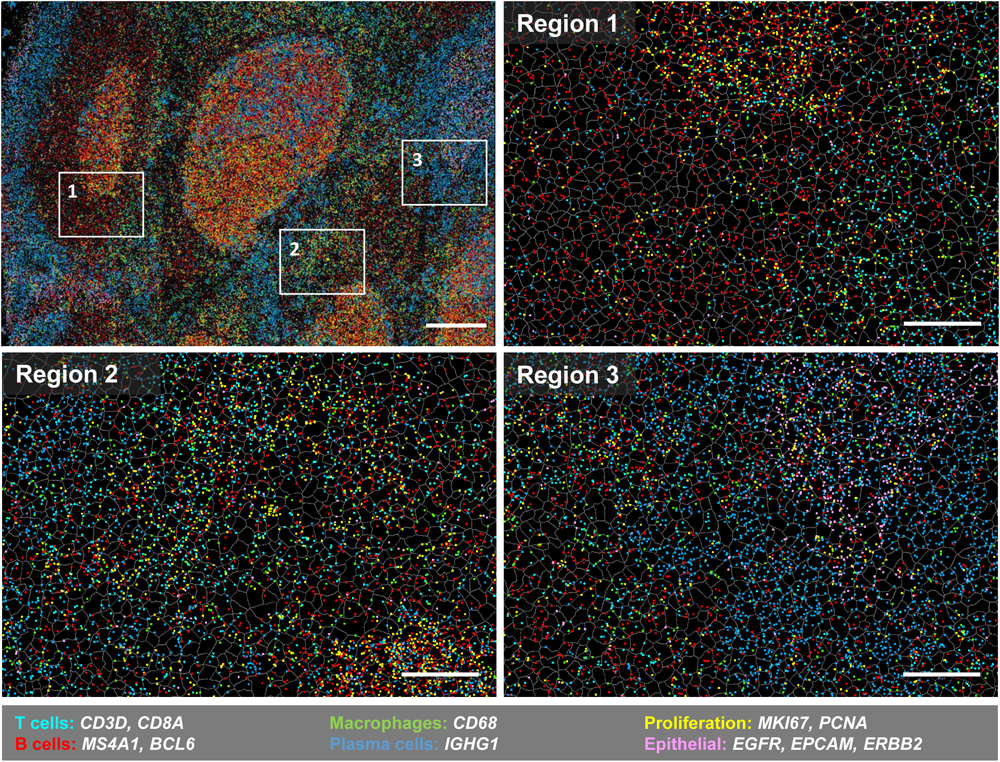
Spatial RNA distribution in normal human tonsil. RNAsky gene expression exhibits histological structure of the tonsil. Numbered boxes on top left image correspond to the three zoomed in images (top right and bottom row). Colored dots indicate location of detected transcripts and grey outlines show cellular segmentation. Scale bar = 200 µm; magnified region scale bar = 50 µm.

Because RNA and protein data were acquired on the same section, the relationship between RNA and protein expression profiles could be assessed within a spatial context (Fig. 3a). For genes such as *CD3D* and *MKI67*, expression was enriched in cells that also expressed the corresponding proteins markers. To identify canonical cell populations of the tonsil based on gene expression profiles, a stringent gating strategy using normalized transcript counts (see methods) was applied to pairwise gene relationships that define different cell types (Fig. 3b, Supplementary Fig. 6a). B cells were defined by the presence of *MS4A1*, while T cells were identified by the absence of this gene and presence of *CD3D* and/or *CD8A*. Then, additional selection gates were applied to further characterize these two types of lymphocytes. Germinal center cells were characterized by double expression of *MS4A1* and *BLC6*, and proliferating cells within this region were further identified with the presence of *MKI67*. Cytotoxic T cells showed a combination of *CD3D* and *CD8A* expression, while remaining T cells were characterized by *CD3D* expression alone. Finally, epithelial cells were defined by the expression of three hallmark genes *EGFR*, *EPCAM*, and *ERBB2* and the absence of *MS4A1*.

**Fig. 3.**
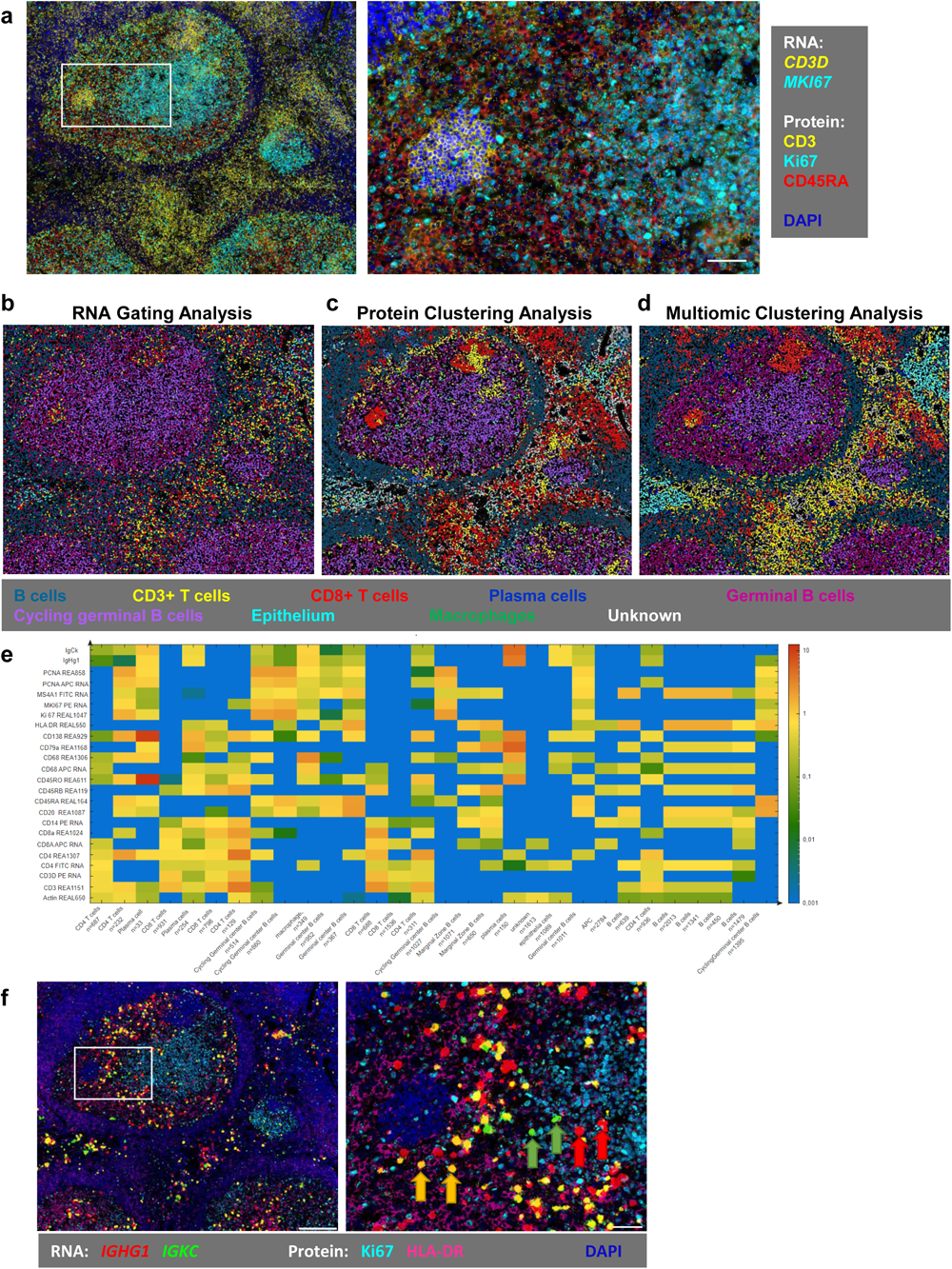
Multiomic analyses by combining spatial information of RNA and protein expression in human tonsil. **a** Display of RNA transcripts with respective proteins on the same section of a human tonsil. Box on left image is location of zoomed in image on right. Colors of protein immunostaining and detected RNAs (dots) indicated. DAPI counterstaining shown in blue. **b** Cell type annotation based on known gene relationships using stringent RNA gating strategy in human tonsil section. Cell types for all analysis strategies (b, c, d) shown below images. **c** Agglomerative hierarchical clustering using protein immunostaining intensity in human tonsil section. **d** Agglomerative hierarchical clustering using protein immunostaining intensity and RNA counts in human tonsil section. **e** Heatmap of selected protein and RNA intensities in spatially grouped, protein-based cell type clusters. The y-axis contains spatially-group clusters, thus there can be more than one cluster of the same cell type, with number of cells in each cluster. **f** Intensity-based transcript detection of IG genes, in a manner similar to antibody-stained tissues, allows for the identification of plasma cells expressing either *IGHG1* mRNA (red arrows), *IGKC* mRNA (green arrows), or both (yellow arrows). Scale bar = 200 µm; close-up region scale bar = 50 µm.

To investigate cell population dynamics of the tonsil, major cell types were determined using hierarchical clustering based on protein signal intensities (Fig. 3c, Supplementary Fig. 6b). Eight major cell types were identified including B cells, T cells, germinal center B cells, proliferating B cells, cytotoxic T cells, plasma cells, macrophages, and epithelial cells. Clustering based on both transcript counts and protein intensities further improved cell population annotation by refining the boundaries between populations (Fig. 3d, Supplementary Fig. 6c). A heatmap of intensity-based quantification of select protein and mRNAs allowed for the visualization of expression differences between spatially distinct cell type clusters (Fig. 3e). For example, one of the five CD4 T cell groups (second column) may be proliferative with higher protein and RNA levels of the proliferation markers PCNA and Ki67. Additionally, this heatmap exhibited correlations of RNA and protein to define cell types, such as plasma cells. This cell type showed a high correlation of *IGHG1* and *IGKC* transcripts, and CD79a and CD138 marker expression. Thus, using the intensity-based detection method, plasma cells were visualized by mRNA expression with cells expressing either *IGHG1* (red arrows), *IGKC* (green arrows), or both (yellow arrows) (Fig. 3f, Supplementary Fig. 7).

### Multiomic analyses capture the dynamics of the tumor microenvironment and expression signatures across different cancer systems

The extended I-O panel enabled robust transcriptional analyses across cancer samples. To determine the panel’s ability to capture canonical cancer and immune signatures, gene expression profiles were assessed across four cancer samples (Fig. 4, Supplementary Fig. 8). High *GATA3* expression is commonly seen in well differentiated breast cancers (BC), and was observed in the triple positive breast cancer sample (Fig.4a) ^40^. The non-keratinizing squamous cell carcinoma (NKSCC) sample showed high *CDKN2A* expression (Fig. 4b). This is in alignment with cBioPortal for Cancer Genomics data sets showing high *CDKN2A* levels in 97% of patients with HPV+ head and neck squamous cell carcinomas ^41, 42, 43^. Immune infiltration is a key factor in understanding cancer progression and prognostic outcomes. Three of the tumor ROIs exhibited the presence of an immune infiltrate. In the breast cancer sample (Fig. 4a), T and B cells can be observed throughout the tumor. The NKSCC (Fig 4b) and colorectal cancer (CRC) (Fig. 4c), showed organization of T and B cells into tertiary lymphoid structures (TLSs), with presence of *BCL6* indicating germinal center formation (Fig. 4c) ^44^.

**Fig. 4.**
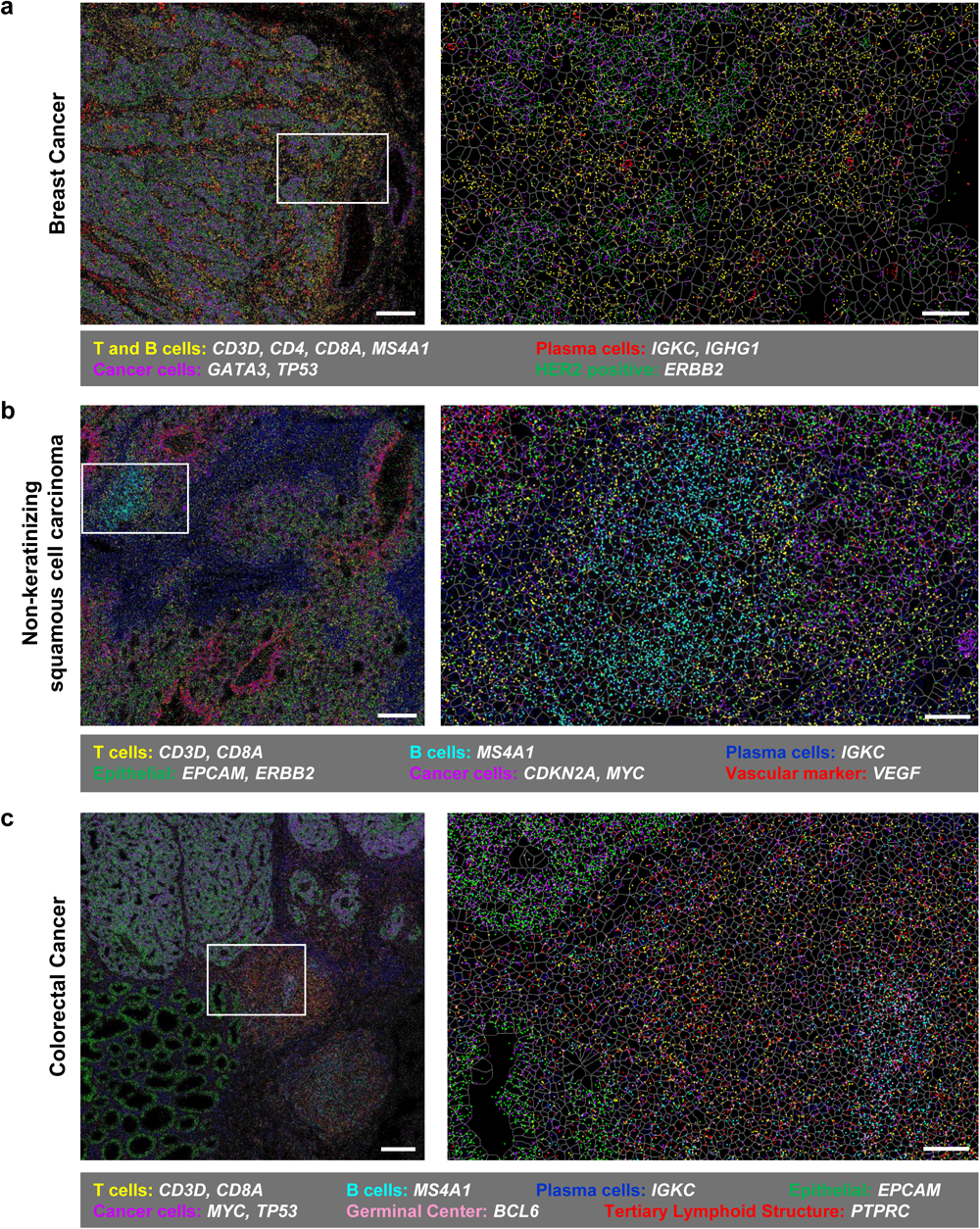
Spatial RNA distribution in multiple different cancer types. RNAsky on a representative **a** breast cancer **b** non-keratinizing squamous cell carcinoma and **c** colorectal cancer. Box on left image is location of zoomed in image on right. Colored dots indicate location of selected transcripts (legend below each image), and grey outlines show cellular segmentation. Scale bar = 200 µm; close-up region scale bar = 50 µm.

The combination of high-resolution transcriptomics and proteomics data enables characterization of the cell interactions and molecular mechanisms occurring within the spatial context of the tumor microenvironment (Fig. 5). A tonsil control was included in these analyses as a normal tissue reference. Each cancer system showed a distinct profile, highlighting the various levels of gene expression in different tumors (Fig. 5a, Supplementary Fig. 9). Following the Seurat pipeline, dimension reduction, clustering, and curated cluster annotation was performed (Fig. 5b, see methods) ^45^. Five cell populations were defined based on gene expression signatures, including cancer/epithelial cells, T cells, B cells, plasma cells, and macrophages/monocytes. (Fig. 5b bottom). Using the cluster annotations, cell population dynamics, both quantitatively and spatially, were evaluated (Fig. 5c-d). The tonsil showed a low presence of the cancer/epithelial cell group, representing the tonsillar crypt epithelial cells, while the tumor ROIs showed a significant amount of malignant epithelial cells. This is reflected in the enrichment of the epithelium population within all tumor ROIs (unpaired t-test with Welch’s correction, p<0.05).

**Fig. 5.**
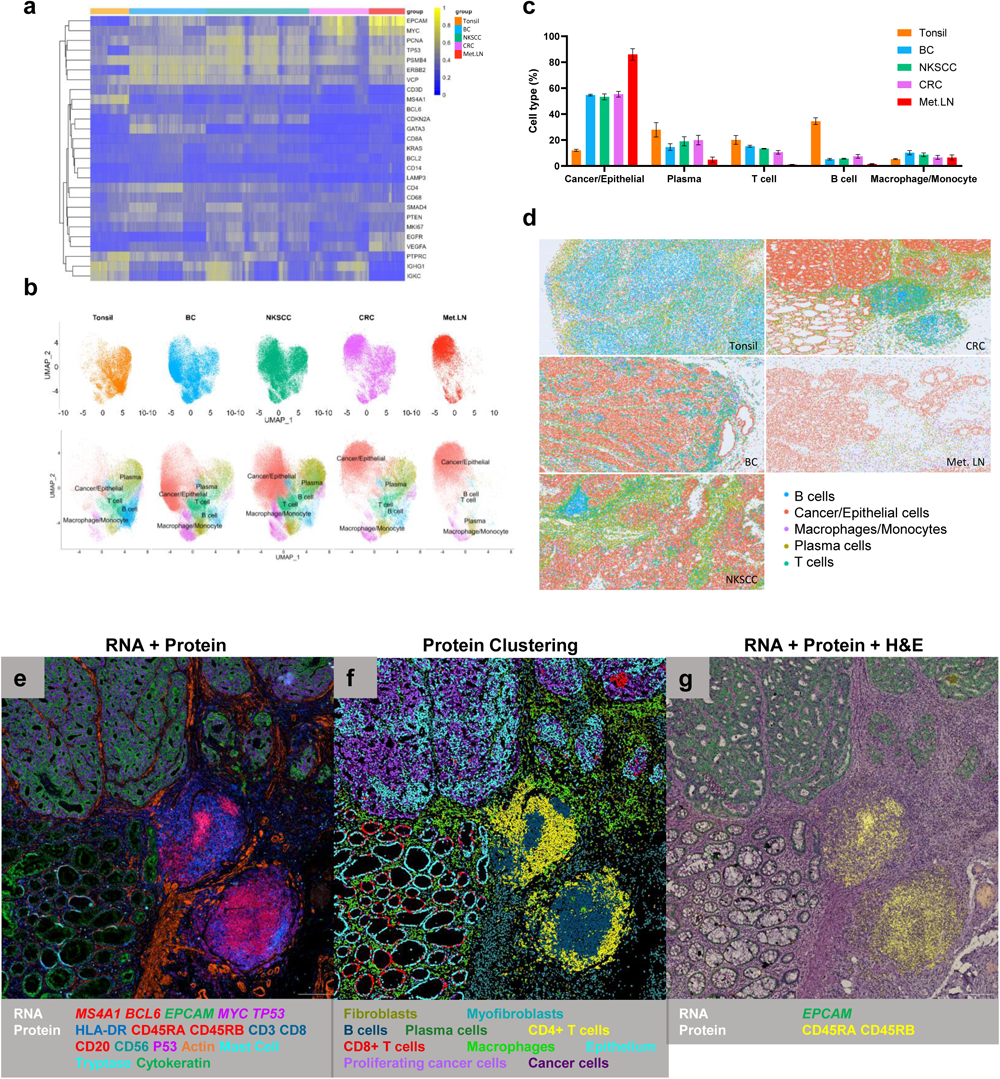
Multiomic analyses across cancer samples. **a** Clustered heat map of normalized gene expression of all genes in the extended I-O panel across all tissue types (BC – breast cancer, NKSCC – non-keratinizing squamous cell carcinoma, CRC – colorectal cancer, Met. LN – metastatic lymph node). **b** UMAP plots of all tissue types analyzed. Annotated cell populations labeled in bottom image. **c** Cross sample quantification of annotated cell populations. Data represents mean ± SEM (n=3 for BC, NKSCC, Met. LN; n=2 for tonsil, CRC). **d** Spatial reconstruction of identified cell populations based on the locations of transcription expression. **e** RNAsky transcript detection and immunostaining exhibit distinct features in colorectal cancer. Colors of detected proteins and RNAs (dots) indicated in legend below image. **f** Agglomerative hierarchical clustering using protein immunostaining intensity in colorectal cancer. Clusters and associated coloring in legend below image. **g** H&E performed on the same tissue section after combined MICS run is registered and combined with DAPI counterstaining (grayscale), then overlaid with RNAsky transcripts and protein immunostaining. Colors of detected proteins and RNAs (dots) indicated in legend below image. Scale bar = 200 µm.

Adding protein information to RNA showed the functional organization of the tumor microenvironment (Fig. 5e, Supplementary Figs. 10-12). The tonsil and nonkeratinizing squamous cell carcinoma showed strong expression for cytokeratin 5 (CK5) and high molecular weight cytokeratin (CK HMW) in the tonsillar crypt epithelium and malignant squamous epithelium respectively (Supplementary Fig. 10). The breast carcinoma sample showed a characteristic cytokeratin 7 (CK7) positive, cytokeratin 20 (CK20) negative cytokeratin profile and the colorectal carcinoma the inverse (CK7 negative / CK20 positive) as expected. As seen in Figure 5e, malignant epithelium (purple and green) was identified in the upper two quadrants of the CRC image, while normal epithelium (green) was located on the bottom left quadrant. Two TLSs (red and blue) were located in the bottom right quadrant of the image, as characterized by their RNA and protein expression and histological appearance. Using hierarchical clustering based on protein expression, ten dominant cell populations were identified representing epithelial cells, cancer cells, proliferating cancer cells, stromal cells, and immune cells (Fig. 5f). Because erasure methods used were gentle, samples remained intact at the completion of image acquisitions, enabling downstream hematoxylin and eosin (H&E) staining on the same tissue section. The H&E images were registered with DAPI images from the RNA and protein image acquisition. Once registered, RNA, protein, and histological data could be simultaneously viewed in the MACS iQ View software platform (Fig. 5g, Supplementary Fig. 13), allowing for direct comparison of classic histology with spatially valuable transcriptomic and proteomic information on the same section.

## Discussion

In this study, we introduced a true multiomics approach, where the chemistry for RNA detection was compatible with subsequent immunofluorescent epitope detection, enabling the combined analysis of RNA and protein targets on a single, histological tissue section. Here, RNAsky showed a high reproducibility for serial sections, which now serves as the basis for spatially unbiased results from MICS technology on the same tissue section. Additionally, because the methods used were gentle, samples remained intact at the completion of all image acquisitions, enabling downstream histology staining (H&E) on the same section. Transcriptomic and proteomic approaches have identified a substantial degree of variation in both, protein and mRNA levels in most, if not all types of cancer ^46, 47, 48^. This observed heterogeneity often increases as tumors progress. In addition, kinetics and intensity of RNA and protein expression of the same gene can differ due to post transcriptional control. For example, miRNAs negatively regulate the expression of a large proportion of mRNAs ^49, 50^ and translational regulation/post translational modifications alter protein stability and abundance ^51, 52^. The visualization of protein and RNA in the context of a canonical diagnostic stain provides the link between mRNA and protein expression and qualitative image analysis/interpretation (H&E). This increases the level of confidence when determining the correlation or the divergence between all three established information modalities. It also enables more accurate phenotyping, determination of cell states, tumor microenvironment profiling, cell segmentation, and clustering, all of which are key to capturing molecular changes contributing to the progression and cellular response of disease. Finally, it can also be used to study the mechanisms of gene regulation in a spatial context.

The approach described here is based on *in situ* hybridization and amplification of probes for RNA transcript detection. The padlock probes in RNAsky were designed to anneal directly to mRNA, eliminating the need for cDNA synthesis therefore improving the detection sensitivity. While we have used a first set of padlocks to evaluate the technology, the addition of new gene targets to a panel is straightforward. As each gene is detected by a single detection probe sequence independently of all other targets, users can readily combine or modify gene panels. Additionally, the size of a panel is flexible and the upper limit is only restricted by the processing time. When we assessed the FDR, calculated across multiple replicates, for the transcript detection approach, the proportion of false positive genes in the extended I-O panel was extremely low (mainly less than <0.1%). This low FDR indicates a high degree of specificity of RNAsky, which requires the hybridization of the two arms of the padlock probes to occur, and a ligation event to generate the amplified molecule underlying the observed punctate signal. Our punctate detection approach shows a higher sensitivity for low and medium expressed genes compared to scRNA-Seq, but a lower sensitivity for highly expressed transcripts. Intuitively, this observation is in alignment with higher transcript expression being more prone to optical and physical overcrowding. To mitigate this issue, we developed a computational approach, integrated into the MACS iQ View analysis software, where the discrete count-based quantification is supplemented with a continuous intensity-based quantification, allowing for a robust quantification and an increase in the dynamic range of detection of RNAsky. Future technological iterations of the MACSima platform will enable z-stack acquisition, where multiple images are taken at different focal distances. By acquiring three-dimensional data, the composite image will have a greater depth of field and will better resolve punctate amplicon signal.

The spatial multiomics analyses performed on the same tissue section revealed the dynamics of cell populations in both healthy and diseased tissues. Distinct profiles of expression and immune signatures based on RNA, protein or a combination could be drawn. Using stringent gating strategies or hierarchical clustering followed by cluster annotation, cell population dynamics were evaluated both quantitatively and spatially. In tonsil, the corresponding mRNA and protein marker expression were enriched in specific cell type cells, allowing the identification of canonical cell populations such as T cells, germinal center B cells, and epithelial cells.

Although scRNA-Seq captures the transcriptional activity of individual cells in tissues, it lacks spatial information. Here, we investigated the spatial expression patterns of RNA and proteins in the microenvironment of tumor. The tumor microenvironment is a complex ecosystem that is composed of various cell types, such as tumor, immune, and stromal cells. This unique environment varies by tumor type and tumor progression over time ^53, 54^. TLS are heterogenous ectopic lymphoid organs that form at inflammatory and cancerous sites. Without the power of spatial immunostaining and transcriptomics information, it is difficult to assess the activation and maturation stage of tumor-associated TLSs. Spatial analyses can identify aggregates of T cells, B cells, and dendritic cells that are characteristic of these structures, and additional protein and transcriptional data can elucidate maturation characteristics.

Given the complexity of studying internal tumor heterogeneity, combining proteomic and transcriptomic data on the same tissue section would increase our understanding of cancer dynamics and biological variations, including the discrepancy observed in mRNA to protein ratio shown recently in multiple cancer types ^46, 47, 48^. The degree to which mRNA levels scale with protein abundances and the implications in cases where this dependency breaks due to dysregulation, could lead to new avenues for cancer research. While we concentrated in our study on the analysis of proteins and mRNAs it is worth noting that the technology can in theory be expanded to other types of molecules. This could include long non-coding RNAs and all kinds of epitopes detectable by antibodies such as cytokines, glycolipids, carbohydrates, or neurotransmitters. The combinatorial approach described here, using the MACSima platform, overcomes a key limitation of current technologies, and could potentially lead to the identification of new clusters or multimodal patterns of expression, further enabling the discovery and development of predictive and prognostic cancer biomarkers.

## Methods

### Biospecimen Samples

PSN1 cell pellet FFPE blocks and normal tonsil FFPE blocks were purchased from AMSBIO (3120-0410, AMS6022 respectively). Cancer blocks were multi-omics grade biospecimens purchased from Indivumed Services.

Colorectal cancer (CRC) – The sample is a low-grade adenocarcinoma of the ascending colon.

Breast cancer (BC) – The sample is an ER+, PR+, HER2 3+ invasive carcinoma of the breast.

Non-keratinizing squamous cell carcinoma (NKSCC) – The sample is an HPV+, poorly differentiated non-keratinizing squamous cell carcinoma of the oropharynx.

Metastatic lymph node (Met. LN) – The sample is a lymph node metastasis of a colorectal adenocarcinoma.

### Tissue Processing

For all tissue blocks, 5 µm sections were acquired on a HM355 S automatic microtome (Epredia), collected on high resolution slides (Miltenyi Biotec), and baked for one hour at 60°C in a symphony™ incubator (VWR) from each of the tissue blocks. The slides were deparaffinized in Histoclear II (National Diagnostics), following the manufacturer’s instructions, and rehydrated in nuclease free water. The slides then underwent antigen retrieval in tris-EDTA citrate (TEC) buffer pH9 for 20 min at 98°C in a PT Module (Epredia). After washing twice with nuclease free water, the slides were mounted into MACSwell Imaging Frames (Miltenyi Biotec).

### Hematoxylin and Eosin (H&E) Staining

H&E stains were performed according to the kit’s instructions (Vector Laboratories, H-3502) for pathological assessment and informed Region of Interest (ROI) selection. After H&E staining, coverslips were mounted with Cytoseal XYL (Epredia). The samples that underwent H&E staining after RNA and protein acquisition were processed in the same method as serial sections.

### RNAsky probes and Immuno-Oncology Core Panel design

RNAsky utilizes targeted padlock probes for gene identification. Each probe contains two arms which hybridize to two adjacent sequences of an mRNA of interest, as well as a gene specific sequence used for detection. The mRNA targeting regions of the probes were designed with Miltenyi Biotec’s proprietary probe design pipeline. The pipeline was optimized to produce specific and sensitive probes by filtering candidates on multiple parameters including melting temperature, GC content, secondary structure formation, homopolymers, repetitive sequences, and off target events. The I-O Core Panel includes 21 gene targets representing canonical and actionable hallmarks across different cancer systems, two housekeeping genes, and three negative control probes. The two housekeeping genes were selected for their expression levels as previously described ^55^, and were used for data normalization. The negative control probes were designed to not bind anywhere in the human transcriptome and provide a measure of specificity of the system. In addition to the I-O Core panel, four custom gene targets were designed and included: *BCL6*, *CD14*, *CD68*, and *LAMP3*. A list of gene targets can be found in Supplementary Table 1.

### RNAsky Sample Preparation

RNAsky (Miltenyi Biotec) sample preparation began immediately after MACSwell Imaging Frame assembly, following Miltenyi’s RNAsky user manual for FFPE sections. In brief, probes were hybridized overnight at 45°C in a humidified chamber. Following hybridization, samples were washed with a stringency buffer, and then probes were ligated overnight at 37°C. After ligation, samples were again washed, and an amplification primer was annealed. Rolling circle amplification (RCA) was performed at 30°C overnight. Finally, samples were crosslinked for one hour at room temperature to stabilize amplicons prior to loading samples on the MACSima system.

### MACSima Imaging Cyclic Staining (MICS) Technology

Cyclic imaging on the MACSima System was performed as previously described ^56, 57, 58^ with the addition of cyclic RNA acquisition to generate multiomic data sets. The MACSima Platform combines widefield microscopy with fully automated liquid handling to perform cyclic imaging. The instrument iteratively stains analytes of interest, acquires multispectral images, and erases the signal at the end of each round. For RNAsky rounds, signal was erased with Miltenyi’s proprietary RNAsky release reagent, while antibody rounds were either optically bleached or signal was removed using the REAlease® Release Reagent (Miltenyi Biotec). Briefly, before loading samples onto the instrument, regions of interest were selected based on a pathologist’s assessment of H&E sections. ROIs were selected by identifying well preserved tissue representative of the anatomical structure and the pathology encountered, including normal adjacent tissue and associated tertiary lymphoid structures when present. Areas of overt necrosis, pigmentation, hemorrhage and artifacts were avoided. The size of ROI for all tonsil samples and metastatic lymph node is 1.6x1.3mm, 2.6x1.8 mm for the NKSCC and breast cancer, and 2.1x2.2 mm for the CRC. A list of gene targets and antibodies can be found in Supplementary Table 2.

### Image Analysis

All datasets contained RNA and protein images and were analyzed in the MACS iQ View analysis software (Miltenyi Biotec). Every field of view was assessed across two exposure times during image preprocessing. Flatfield and distortion corrections were applied based on calibration data from the instrument ^33^. In the event of pixel overexposure with the highest exposure time, the signal was normalized with the lower exposed image to generate a high dynamic range image (HDR). Then, DAPI images were used for stitching and registration across the entire ROI for each cycle. Finally, signal images had the background subtracted based on background images acquired for every cycle. Segmentation was performed based on DAPI images in a two-step process, with nuclear segmentation based on the watershed method followed by a donut based cytoplasmic segmentation. A gaussian smoothing was performed on DAPI images to improve cell segmentation for tumor tissues. After segmentation cell features are measured across all channels.

#### Transcript Detection

Utilizing the integrated algorithm in MACS iQ View, transcripts were detected with default parameters. With this algorithm, transcripts are detected in two stages for enhanced accuracy. First, a multi-scale object detection strategy is employed utilizing the Laplacian of Gaussian (LoG) filter. This process evaluates the maximum response over a range of Gaussian kernel sizes, allowing for the identification of objects of various sizes and across multiple focal planes. The outcome of this process is an image characterized by the Laplacian transformation, which is then thresholded to create a collection of high-response regions where punctate signal has been detected. In the second stage, in order to obtain an exact list of object positions, wavelet filtering of the preprocessed images is performed followed by local peak detection. Any local maxima not found within the high-response regions from stage 1 are discarded as false positives, while the remainder form our final curated set of detected transcripts.

#### RNA gating cell population analysis

After transcript detection, gene counts were normalized using the MinMax algorithm in MACS iQ View. Briefly, the MinMax normalization algorithm transforms all values for a given feature into the range [0,1], where the minimum and maximum value for that feature is 0 and 1 respectively. Then, pairwise scatterplots were generated, and selection gates were established based on desired gene relationships. For example, when using gene A and gene B, a user can select a gate for double positive expression (A and B), or an exclusion gate (A not B). A very stringent approach was used where cells were not allowed to be represented in more than one population group.

#### Protein clustering analysis

For protein clustering, the fluorescence intensity of each marker was measured and a Z-normalization used across all protein measurements. Then, principal component analysis for 4 dimensions was performed. These were then used as the input for the hierarchical clustering within MACS iQ View with the following parameters: agglomerative hierarchical clustering algorithm, number of clusters 30, Euclidean distance (L2 norm), non-squared, and Ward’s method.30 clusters were generated, and cell types were assigned to a cluster based on the five most expressed markers. Clusters were combined when the same cell type was assigned in separate clusters. The identified cell types were colored and plotted in an XY plot using the nuclear segmentation mask as the underlying map. Expression heatmaps were generated displaying both protein and RNA intensity values across the annotated spatially grouped protein-based cell type clusters.

#### Multiomic clustering analysis

RNA count data and protein intensity data were used as input features for the agglomerative hierarchical clustering algorithm in MACS iQ View with the following parameters: number of clusters 20, Euclidean distance (L2 norm), non-squared, and Ward’s method. After initial clustering, clusters were filtered and annotated to show cell types.

#### Intensity-based Transcript Quantification

The overall intensity of signals from detected RNA transcripts in segmented cells was measured in the same way the amount of signal from fluorescent antibodies is assessed. The intensity-based method is compatible with software developed for fluorescence microscopy and uses the same methodology as the antibody staining method. For the comparison of both count-and intensity-based transcript quantification, the DAPI image and the RNAsky image of the *MS4A1* (CD20) were imported into QuPath ^59^. The cell nuclei were segmented using the DAPI channel and a defined area around the nuclei was used as the cytoplasm. Using the experimental organelle detection method within QuPath, the detected RNA transcripts were segmented and counted per cell. From these data both the intensity and the count data were extracted and exported into a data table and further analyzed.

#### MICS and H&E registration

H&E staining was performed following RNAsky and MICS run on MACSima as described above, and whole slide images were acquired. For image registration of H&E image to DAPI channel, selected ROI area was cropped and then stacked with inverted cycle 0 DAPI image in Fiji. The stacked images were aligned using Fiji’s plugin, Linear Stack Alignment with SIFT^60^ and merged by Z projection in Fiji. To combine transcripts, protein and H&E images, the registered H&E image was imported into MACS iQ View analysis software. The stacked images were aligned using Fiji’s plugin, Linear Stack Alignment with SIFT ^60^ and merged by Z projection in Fiji. To combine transcripts, protein and H&E images, the registered H&E image was imported into MACS iQ View and overlaid with detected transcripts and protein images acquired from MACSima.

### RNAsky performance and cross-technology comparisons

For the following analyses, *IGHG1*, *IGKC* counts were excluded due to their very high expression.

#### False Discovery Rate (FDR) Calculation

FDR (%), a measure of specificity, is the rate of calls from negative control probes (n=3) per total number of calls for a region of interest (ROI). The FDR calculation was determined based on the I-O Core Panel for Tonsils 1 and 2 and the extended I-O Panel for all remaining tissues.

#### Bulk Sequencing Correlation

Pearson correlation with 95% confidence intervals was determined on the I-O Core Panel genes between the mean counts per ROI of a representative tonsil sample and nTPM of a bulk RNA-Seq tonsil dataset. Two-tailed P value was also determined.

#### Cell Line Correlation

The counts or TPM for each gene were normalized to *PSMB4* counts or TPM. Pearson correlation with 95% confidence intervals was determined on the I-O Core Panel genes between the average of two PSN1 RNAsky normalized counts and a PSN1 Cancer Cell Line Encyclopedia (CCLE) bulk RNA-Seq dataset. Two-tailed P value was also determined.

#### Single Cell Sequencing (scRNA-Seq) Correlation

Pearson correlation with 95% confidence intervals was determined on the I-O Core Panel genes between the mean counts per cell of a representative tonsil sample and mean counts per cell of a scRNA-Seq healthy young adult tonsil dataset. Two-tailed P value was also determined.

#### In situ Sensitivity Calculations

Both RNAsky and scRNA-Seq are able to measure transcripts per cell allowing for the determination of how sensitive RNAsky *in situ* transcript detection is relative to scRNA-Seq transcript detection. In healthy tonsil RNAsky datasets and a scRNA-Seq healthy young adult tonsil dataset, transcripts/cell was determined for each gene followed by the ratio of RNAsky to scRNA-Seq for each gene. Median sensitivity was determined from the ratios all the I-O Core Panel genes.

### Gene expression analyses

#### Transcript count preprocessing: filtering and transcript count normalization

For each sample, cells with less than three RNAsky genes detected were filtered out, and RNAsky genes detected in less than three cells were excluded from downstream analysis. The raw transcript counts from all RNAsky samples were first merged to a single Seurat object by Seurat (version 4.3.0) function ‘merge’ ^45^. To normalize transcript counts across all cells, the raw gene transcript count for each cell was divided by the total transcript count of that cell, multiplied by scale factor 10,000, and then natural log transformed with pseudo count set as 1. Seurat function NormalizeData(normalization.method=”LogNormalize”, scale.factor=1e4) was used for the log normalization. Subsequently, the log normalized counts were transformed to Z-score by ‘Seurat::ScaleData()’, and finally gene expression variance was stabilized by ‘Seurat::SCTransform’.

#### RNAsky transcript count dimensionality reduction and batch effect correction

Principle Component Analysis was performed on the merged object as described above with all RNAsky genes included and total number of PCs set as 26 (Seurat::RunPCA(npcs=26)). To correct batch effect, all samples in the merged object were integrated by the fast, sensitive, and scalable integration tool Harmony, and performed by Seurat wrapper ‘RunHarmony’ ^61, 62^. Cells from individual samples were considered as different batches (specified by RunHarmony parameter group.by). The integration process generated harmony components, which were all used subsequently as input for UMAP (Seurat::RunUMAP(dims=1:26)) and clustering analysis.

#### Clustered heatmap plotting, cell type annotation and spatial reconstruction of cell types

The single cell clustered heatmap was generated based on the above log normalized counts, which was scaled between 0 and1. The plotting was performed by the R function dittoSeq::dittoHeatmap(scaled.to.max=TRUE).

To cluster cells for cell type annotation, all the harmony components were utilized to construct a shared nearest neighbor (SNN) graph. This was performed by Seurat::FindNeighbors(dims=1:26) function with the other parameters kept at their default values. Subsequently Louvain algorithm was employed to identify clusters by Seurat::FindClusters(algorithm=1, resolution=0.1). The “resolution” parameter for Louvain clustering determines the size and number of detected clusters. In this study, we tested resolution values ranging from 0.1 to 0.5. When resolution was set at 0.1, a total of seven clusters were identified, which aligns well with the expected number of cell types.

The resulting clusters were automatically annotated by R package SingleR (version 2.0.0) against two general-purpose built-in references: Human Primary Cell Atlas and Blueprint/ENCODE,which are provided by R package celldex (version 1.8.0) ^63^. Additionally, we annotated the clusters by R package Azimuth (version 0.4.6) function ‘RunAzimuth’ against the Human tonsil reference ^45^. Marker genes for each cluster were identified by Seurat::FindAllMarkers(test.use = “wilcox”, min.pct=0.1) function with log normalized counts as input and the default Wilcoxon rank sum test. The average of log normalized counts for each cluster was visualized by R function ‘pheatmap::pheatmap’ (version 1.0.12). The final annotation was determined by considering both automatic cell annotation and marker gene expression level.

The spatial reconstruction of cell types was achieved by the ‘scatter’ method in Python library plotly.express (Plotly Technologies Inc.). The input data is the cell coordinates generated by cell segmentation.

### Datasets

The gene expression for all 23 targets in the RNAsky Immuno-Oncology Core Panel was retrieved from the following data sources:

#### Human Protein Atlas (HPA) Healthy Tonsil RNASeq

The RNA HPA tissue gene data set was downloaded from the Human Protein Atlas (http://www.proteinatlas.org) and is based on the Human Protein Atlas version 23.0 and Ensembl version 109 ^8, 64^.

#### Single Cell Tonsil Data Set

Single cell RNA sequencing data was downloaded from the deposited Seurat objects as described in the tonsil atlas publication ^65^. The non-cell-hashed dataset from three healthy young adult donors ages 26, 33, and 35 were used for the analysis in this study. Single cell RNA sequencing data was downloaded from the deposited Seurat objects as described in the tonsil atlas publication ^65^.

#### Cancer Cell Line Encyclopedia (CCLE) RNASeq

The CCLE data set was downloaded from the DepMap Public 22Q2 release on the DepMap portal (http://depmap.org). The PSN1 cell line was selected as a representative cancer system for marker assessment ^66^.

### Statistics

Unless otherwise specified, all bar graphs show the mean with error bars representing the standard error of the mean. Pearson correlation coefficients with 95% confidence intervals and two-tailed P values were determined for all correlations. Unpaired t tests with Welch’s correction with two-tailed P values were performed for comparisons. Statistical analysis performed using GraphPad Prism 10.

## Acknowledgements

We thank Dr. Christian Wöhle for bioinformatics support. We thank Rami Hayajneh for histology image acquisition support. We thank the Molecular Chemistry team in Waltham, MA for their support in RNAsky reagent and protocol developments. This publication is part of the Human Cell Atlas – www.humancellatlas.org/publications/.

## Author contributions

E.N., D.P., R.C.H., R.P., W.M., T.W., A.B, J.F., S.R. designed studies.

E.N., D.P., R.C.H., M.D., E.D., H.L., E.L., H.L., S.S. performed experiments.

E.N., D.P., R.C.H., R.P., J.W., W.M., T.W., A.B., A.M. wrote manuscript.

E.N., D.P., R.C.H., J.W., W.M., T.W., A.M., F.S., D.M., P.P., H.L., E.L., J.F. analyzed the data and interpreted results

E.N., D.P., R.C.H., J.W., W.M., T.W., R.P., A.B., J.F., H.M., T.R., R.H. revised and edited the manuscript

## Materials & Correspondence

Correspondence and material requests should be addressed to Robert Pinard or Andreas Bosio.

## Data availability

As examples, the RNAsky and antibody-stained original stitched images of the Tonsil and the CRC tissue shown in Figures 4 and 5, are uploaded to Zenodo in the OME.TIF file format and the links will be provided after acceptance of the manuscript.

## Supplementary Information

In attached Supplementary Information file.

