## Supplemental Information for "Spatial protein and RNA analysis on the same tissue section using MICS technology"

### Supplementary Information

#### Supplementary Tables


Supplementary Table 1. RNAsky gene targets in extended I-O panel.

Supplementary Table 2. Protein targets and associated antibodies used.


#### Supplementary Figures


Supplementary Fig. 1. Performance metrics of RNAsky.

Supplementary Fig. 2. Count-based and intensity-based methods for RNA quantification in human tonsil.

Supplementary Fig. 3. Spatial distribution of genes in region 1 of a healthy human tonsil.

Supplementary Fig. 4. Spatial distribution of genes in region 2 of a healthy human tonsil.

Supplementary Fig. 5. Spatial distribution of genes in region 3 of a healthy human tonsil.

Supplementary Fig. 6. Analysis approaches for multiomic data.

Supplementary Fig. 7. Intensity-based quantification of *IGHG1* and *IGKC* allows for detection of plasma cells expressing different antibody heavy and light chain combinations.

Supplementary Fig. 8. Spatial RNA distribution in a metastatic lymph node.

Supplementary Fig. 9. Gene expression profiles across cancer samples.

Supplementary Fig. 10. Immunostaining of cytokeratins for pathological tissue assessment.

Supplementary Fig. 11. Spatial distribution of individual genes in a colorectal cancer.

Supplementary Fig. 12. Immunostaining of individual proteins in a colorectal cancer.

Supplementary Fig. 13. Registration of H&E image to DAPI channel.

**Supplementary Table 1**. RNAsky gene targets in extended I-O panel.

| Detected Genes | RefSeq Gene ID | RefSeq or Ensembl Transcript ID |
|---|---|---|
| *BCL2* | 596 | NM_000633.3 |
| *BCL6* | 604 | NM_001706.5 |
| *CD14* | 929 | NM_000591.4 |
| *CD3D* | 915 | NM_000732.6 |
| *CD4* | 920 | NM_000616.5 |
| *CD68* | 968 | NM_001251.3 |
| *CD8A* | 925 | NM_001768.7 |
| *CDKN2A* | 1029 | NM_000077.5 |
| *EGFR* | 1956 | NM_005228.5 |
| *EPCAM* | 4072 | NM_002354.3 |
| *ERBB2* | 2064 | NM_004448.4 |
| *GATA3* | 2625 | NM_001002295.2 |
| *IGHG1* | 3500 | ENST00000631466.1 |
| *IGKC* | 3514 | ENST00000390237.2 |
| *KRAS* | 3845 | NM_004985.5 |
| *LAMP3* | 27074 | NM_014398.4 |
| *MKI67* | 4288 | NM_002417.5 |
| *MS4A1* | 931 | NM_152866.3 |
| *MYC* | 4609 | NM_002467.6 |
| *PCNA* | 5111 | NM_182649.2 |
| *PSMB4* | 5692 | NM_002796.3 |
| *PTEN* | 5728 | NM_000314.8 |
| *PTPRC* | 5788 | NM_002838.5 |
| *SMAD4* | 4089 | NM_005359.6 |
| *TP53* | 7157 | NM_000546.6 |
| *VCP* | 7415 | NM_007126.5 |
| *VEGF* | 7422 | NM_001171623.1 |

**Supplementary Table 2**. Protein targets and associated antibodies used.

| Target name | Uniprot accession number | Gene Symbol | HGNC ID | Host | Isotype | Clone | Vendor | Catalog number | RRID |
|---|---|---|---|---|---|---|---|---|---|
| Actin (smooth muscle) | P62736 | ACTA2 | 130 | human | IgG1 | REAL650 | Miltenyi Biotec | 130-123-363 | AB_2857589 |
| beta Catenin | P35222 | CTNNB1 | 2514 | human | IgG1 | REA480 | Miltenyi Biotec | 130-123-546 | AB_2819492 |
| CD138, Syndecan | P18827 | SDC1 | 10658 | human | IgG1 | REA929 | Miltenyi Biotec | 130-115-479 | AB_2857403 |
| CD20 Cytoplasmic | P11836 | MS4A1 | 7315 | human | IgG1 | REA1087 | Miltenyi Biotec | 130-118-292 | AB_2857425 |
| CD326 | P16422 | EPCAM | 11529 | human | IgG1 | REA1311 | Miltenyi Biotec | 130-127-909 | AB_2928416 |
| CD3e | P07766 | CD3E | 1674 | human | IgG1 | REA1151 | Miltenyi Biotec | 130-120-267 | AB_2876969 |
| CD4 | P01730 | CD4 | 1678 | human | IgG1 | REA1307 | Miltenyi Biotec | 130-127-906 | AB_2921815 |
| CD45RA | P08575 | PTPRC | 9666 | human | IgG1 | REAL164 | Miltenyi Biotec | 130-112-097 | AB_2819365 |
| CD45RB | P08575 | PTPRC | 9666 | human | IgG1 | REA119 | Miltenyi Biotec | 130-126-373 | AB_2876979 |
| CD45RO | P08575 | PTPRC | 9666 | human | IgG1 | REA611 | Miltenyi Biotec | 130-120-095 | AB_2857502 |
| CD56 | P13591 | NCAM1 | 7656 | human | IgG1 | REAL1142 | Miltenyi Biotec | 130-128-467 | AB_2905078 |
| CD66b | P31997 | CEACAM8 | 1820 | human | IgG1 | REA306 | Miltenyi Biotec | 130-122-922 | AB_2811406 |
| CD68 | P34810 | CD68 | 1693 | human | IgG1 | REA1306 | Miltenyi Biotec | 130-128-345 | AB_2905104 |
| CD79a | P11912 | CD79A | 1698 | human | IgG1 | REA1168 | Miltenyi Biotec | 130-120-933 | AB_2784279 |
| CD8a | P01732 | CD8A | 1706 | human | IgG1 | REA1024 | Miltenyi Biotec | 130-117-200 | AB_2857414 |
| Cytokeratin 20 | P35900 | KRT20 | 20412 | human | IgG1 | REAL753 | Miltenyi Biotec | 130-125-685 | AB_2877036 |
| Cytokeratin 5 6 8 17 19 | P13647, P02538, P04259, P48668, P05787, Q04695, P08727 | KRT5, KRT6A, KRT6B, KRT6C, KRT8, KRT17, KRT19 | 6442, 6443, 6444, 20406, 6446, 6427, 6436 | human | IgG1 | REA1141 | Miltenyi Biotec | 130-120-096 | AB_2857503 |
| Cytokeratin 5 | P13647 | KRT5 | 6442 | human | IgG1 | REAL899 | Miltenyi Biotec | 130-127-016 | AB_2905179 |
| Cytokeratin 7 | P08729 | KRT7 | 6445 | human | IgG1 | REA935 | Miltenyi Biotec | 130-115-448 | AB_2857395 |
| Cytokeratin HMW | P04264, P13647, P13645, P02533 | KRT1, KRT5, KRT10, KRT14 | 6412, 6442, 6413, 6416 | human | IgG1 | REAL645 | Miltenyi Biotec | 130-125-779 | AB_2877037 |
| ErbB2 | P04626 | ERBB2 | 3430 | human | IgG1 | REA1315 | Miltenyi Biotec | 130-127-979 | AB_2905194 |
| HLA DR | P01903, P01911, P79483, P13762, Q30154 | HLA-DRA, HLA-DRB1, HLA-DRB3, HLA-DRB4, HLA-DRB5 | 4947, 4948, 4951, 4952, 4953 | human | IgG1 | REAL550 | Miltenyi Biotec | 130-123-076 | AB_2857572 |
| Ki67 | P46013 | MKI67 | 7107 | human | IgG1 | REAL1047 | Miltenyi Biotec | 130-127-837 | AB_2905309 |
| Mast Cell Tryptase | Q8IX19 | MCEMP1 | 27291 | human | IgG1 | REAL798 | Miltenyi Biotec | 130-125-278 | AB_2857771 |
| p53 | P04637 | TP53 | 11998 | human | IgG1 | REA1132 | Miltenyi Biotec | 130-119-502 | AB_2857470 |
| PCNA | P12004 | PCNA | 8729 | human | IgG1 | REA858 | Miltenyi Biotec | 130-114-702 | AB_2726763 |
| Podoplanin | Q86YL7 | PDPN | 29602 | human | IgG1 | REA446 | Miltenyi Biotec | 130-128-776 | AB_2921938 |

a

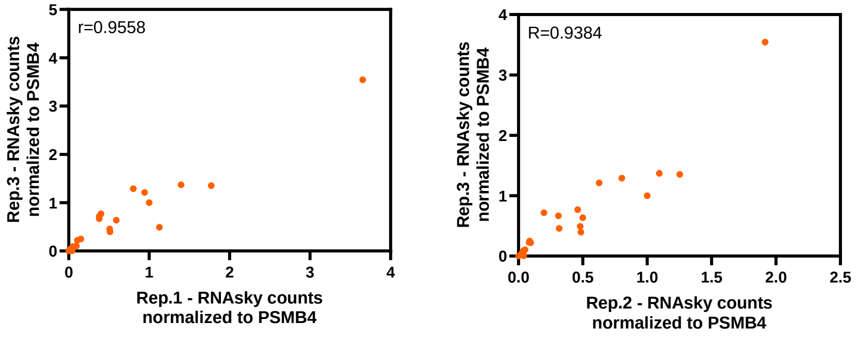


b

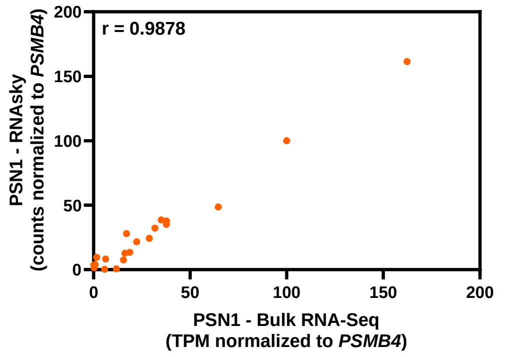


c

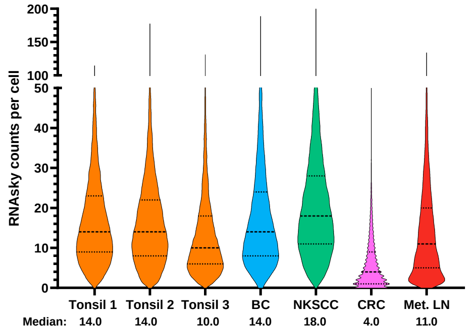


| Total cell numbers | | |
|---|---|---|
| Tissue | Mean | SD |
| Tonsil 1 | 22782 | 977 |
| Tonsil 2 | 21313 | 1131 |
| Tonsil 3 | 26222 | 1155 |
| BC | 31237 | 1074 |
| NKSCC | 44267 | 2473 |
| CRC | 49323 | 293 |
| Met. LN | 14768 | 1598 |

d

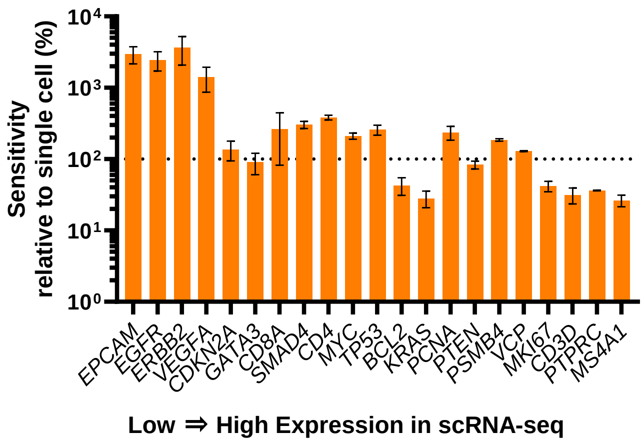


**Supplementary Fig. 1. Performance metrics of RNAsky.**
**a** Correlation of RNAsky counts between independent runs (replicates) of a representative healthy tonsil tissue. Rep. 1 vs Rep. 3 Pearson correlation coefficient (PCC) = 0.9558 (95% confidence intervals = 0.8947 to 0.9818); P value (two-tailed) <0.0001. Rep. 2 vs Rep. 3 PCC = 0.9384 (95% confidence intervals = 0.8551 to 0.9745); P value (two-tailed) <0.0001. **b** Correlation of PSN1 cell line RNA-Seq dataset (Cancer Cell Line Encyclopedia (CCLE) RNASeq) and RNAsky counts of a PSN1 cell line plug (n = 2). PCC = 0.9878 (95% confidence intervals = 0.9696 to 0.9951); P value (two-tailed) <0.0001. **c** Left, Range of total transcripts per cell in representative example of each tissue (BC – breast cancer, NKSCC – non-keratinizing squamous cell carcinoma, CRC – colorectal cancer, Met. LN – metastatic lymph node). Median and quartiles shown by dashed lines. Right, Average number of cells and standard deviation of all tissues analyzed. **d** Average per gene *in situ* sensitivity of RNAsky in healthy tonsils relative to a healthy young adult tonsillar scRNA-seq dataset. Genes ordered from low to high transcripts per cell (expression) in the scRNA-seq data set. The line at 100% indicates the ratio where the sensitivity of scRNA-Seq and RNAsky are equal. Data represents mean ± SEM (n=3 tonsils).

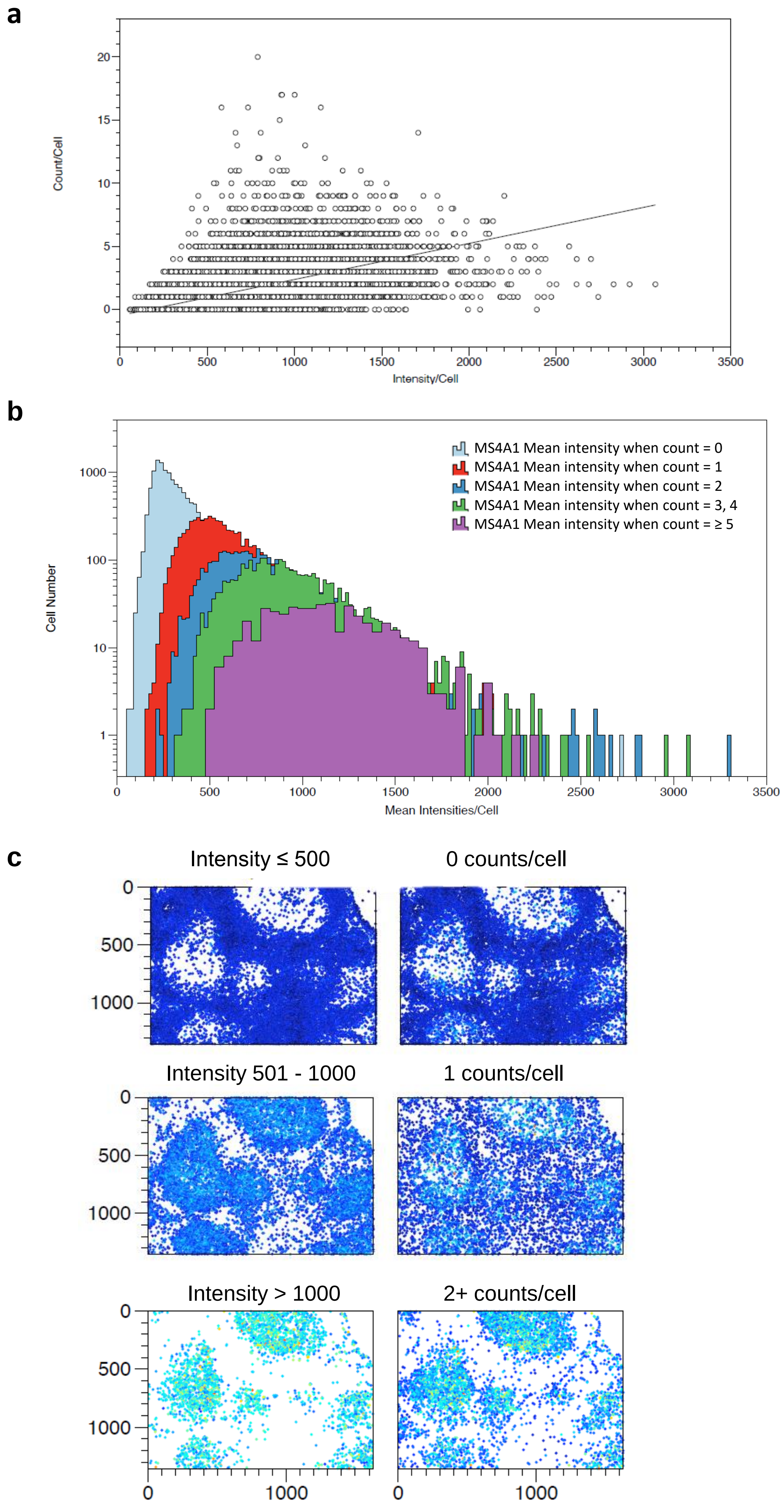
a
Count/Cell
Intensity/Cell
b
MS4A1 Mean intensity when count = 0
MS4A1 Mean intensity when count = 1
MS4A1 Mean intensity when count = 2
MS4A1 Mean intensity when count = 3, 4
MS4A1 Mean intensity when count = ≥ 5
Cell Number
Mean Intensities/Cell
c
Intensity ≤ 500
0 counts/cell
Intensity 501 - 1000
1 counts/cell
Intensity > 1000
2+ counts/cell

**Supplementary Fig. 2. Count-based and intensity-based methods for RNA quantification in human tonsil.**

**a** Correlation between count and intensity of RNA transcript detection measurements for *MS4A1*. Intensity data is continuous, while the count data is discrete. **b** Intensity histograms gated on counts per cell. Cells were gated on the number of *MS4A1* RNA transcripts detected per cell and then the intensity of each gated cell population was plotted as a histogram. The correlation between count/cell and mean intensity/cell is visible with the histograms shifting to the right with increasing counts/cell. **c** Comparison between intensity- and count-based gating showing that both gating strategies work and give similar results. The first row shows cells with an intensity level of *MS4A1* transcripts up to 500 (left) and 0 counts/cell of *MS4A1* (right). The second row shows cells with an intensity level of *MS4A1* transcripts between 501 and 1000 (left) and 1 count/per cell of *MS4A1* (right). The third row shows cell with an intensity level of *MS4A1* transcripts above 1000 (left) and 2 or more counts/cell of *MS4A1* (right).

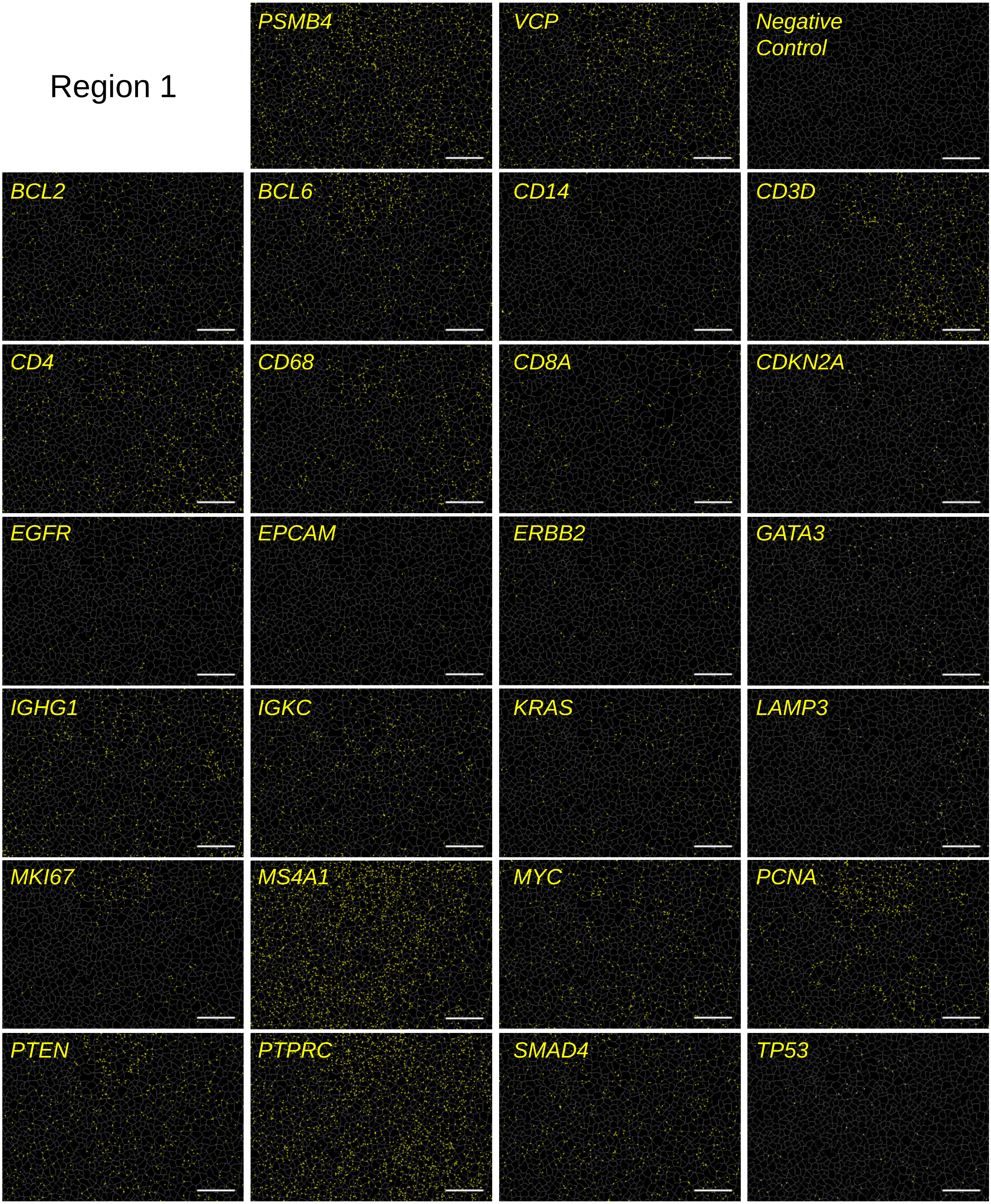


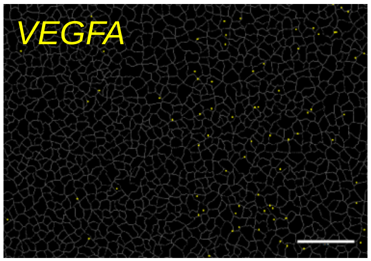


**Supplementary Fig. 3. Spatial distribution of genes in region 1 of a healthy human tonsil.**
Location of RNA transcripts detected by RNAsky for each of the genes in the extended I-O panel set from Region 1 in Fig. 2. Housekeeping genes (*PSMB4* and *VCP*) and negative control in top row. Scale bars = 50 µm.

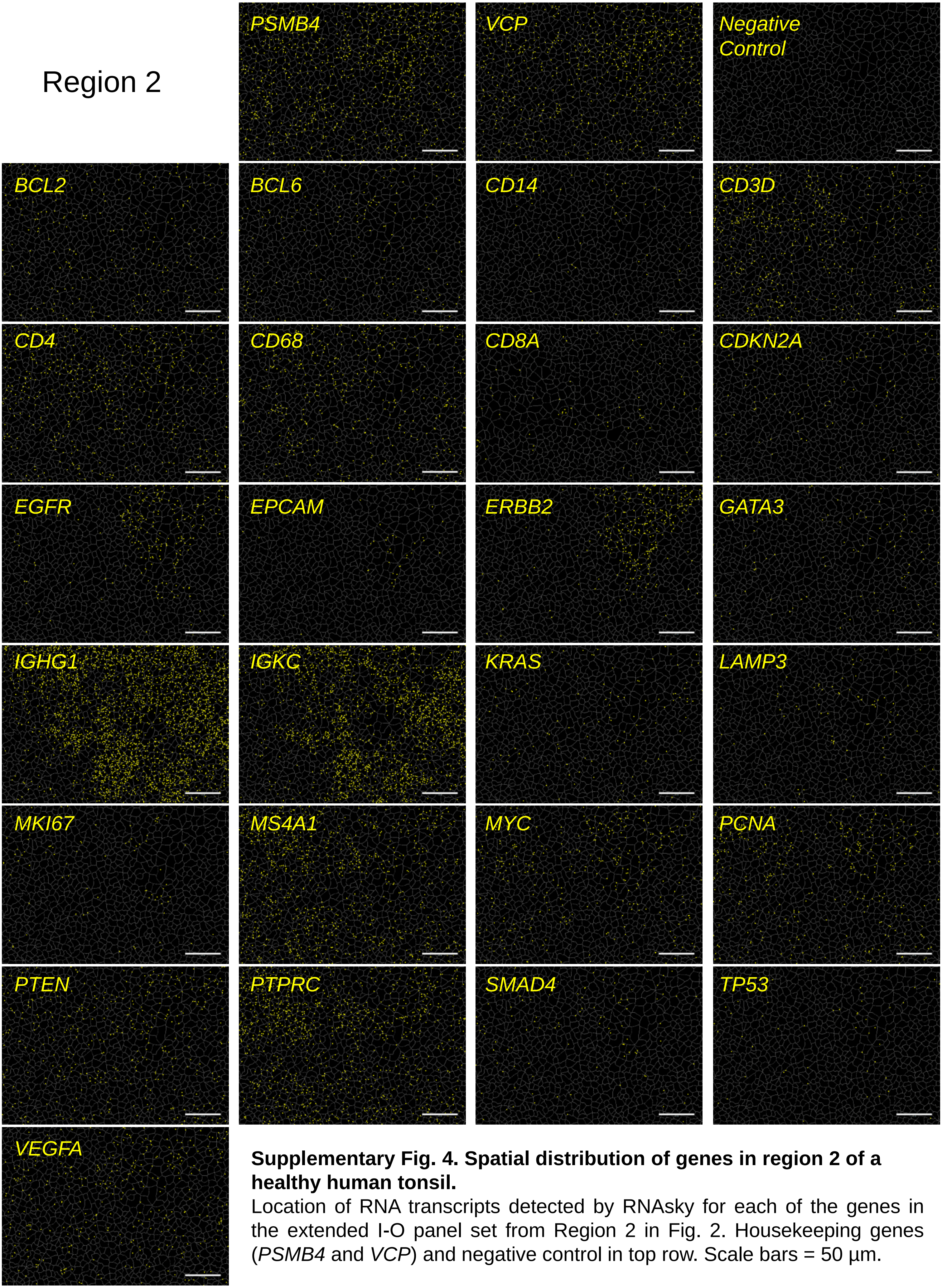


**Supplementary Fig. 4. Spatial distribution of genes in region 2 of a healthy human tonsil.**
Location of RNA transcripts detected by RNAsky for each of the genes in the extended I-O panel set from Region 2 in Fig. 2. Housekeeping genes (*PSMB4* and *VCP*) and negative control in top row. Scale bars = 50 µm.

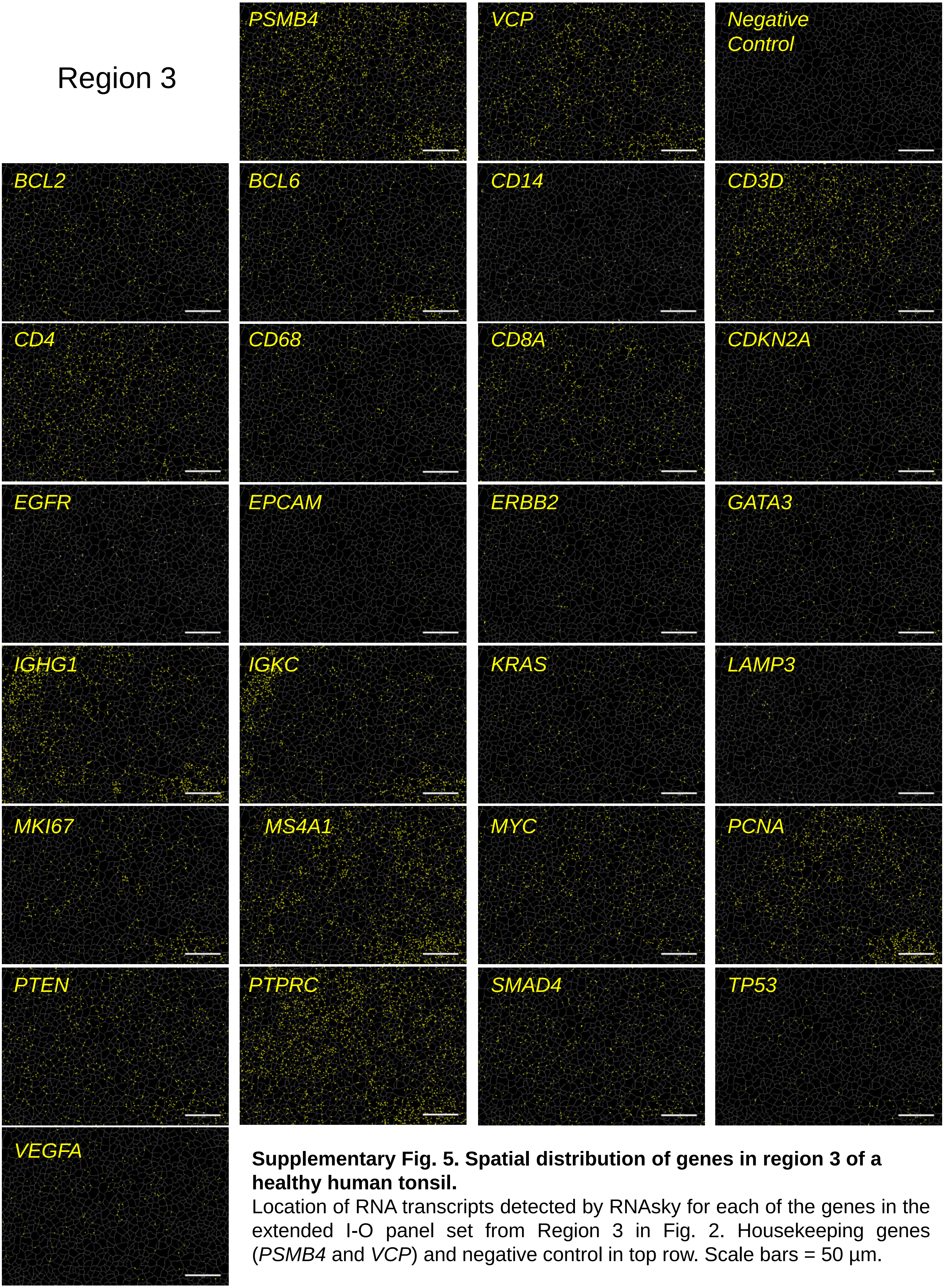


**Supplementary Fig. 5. Spatial distribution of genes in region 3 of a healthy human tonsil.**
Location of RNA transcripts detected by RNAsky for each of the genes in the extended I-O panel set from Region 3 in Fig. 2. Housekeeping genes (*PSMB4* and *VCP*) and negative control in top row. Scale bars = 50 µm.

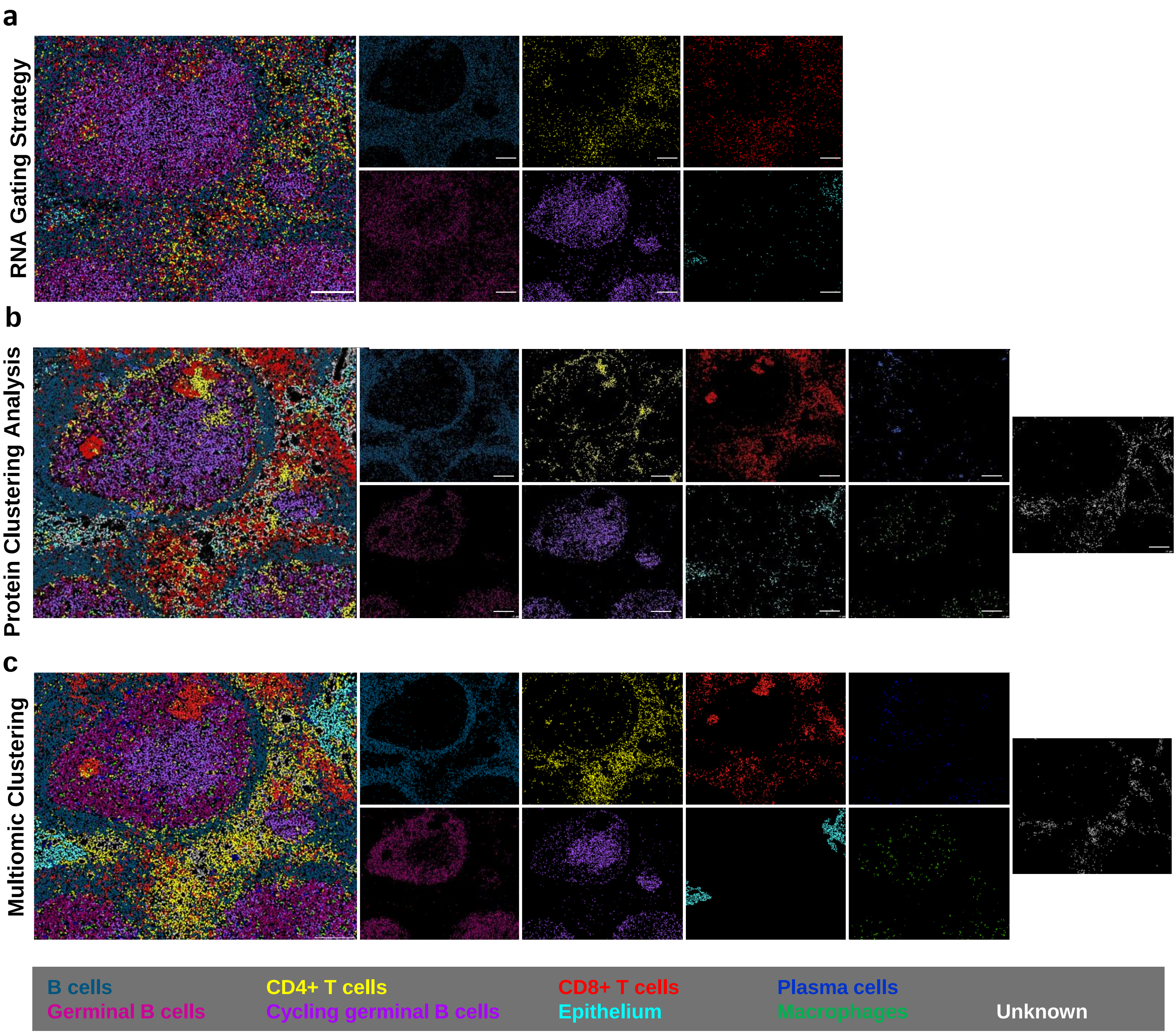


**Supplementary Fig. 6. Analysis approaches for multiomic data.**
**a, b, c** Individual images of identified cell groups/populations after clustering. a, b, c show results of clustering methods exhibited in Fig 3b, d and f, respectively. Clusters and coloring shown at bottom. Image scale bars = 200 µm.

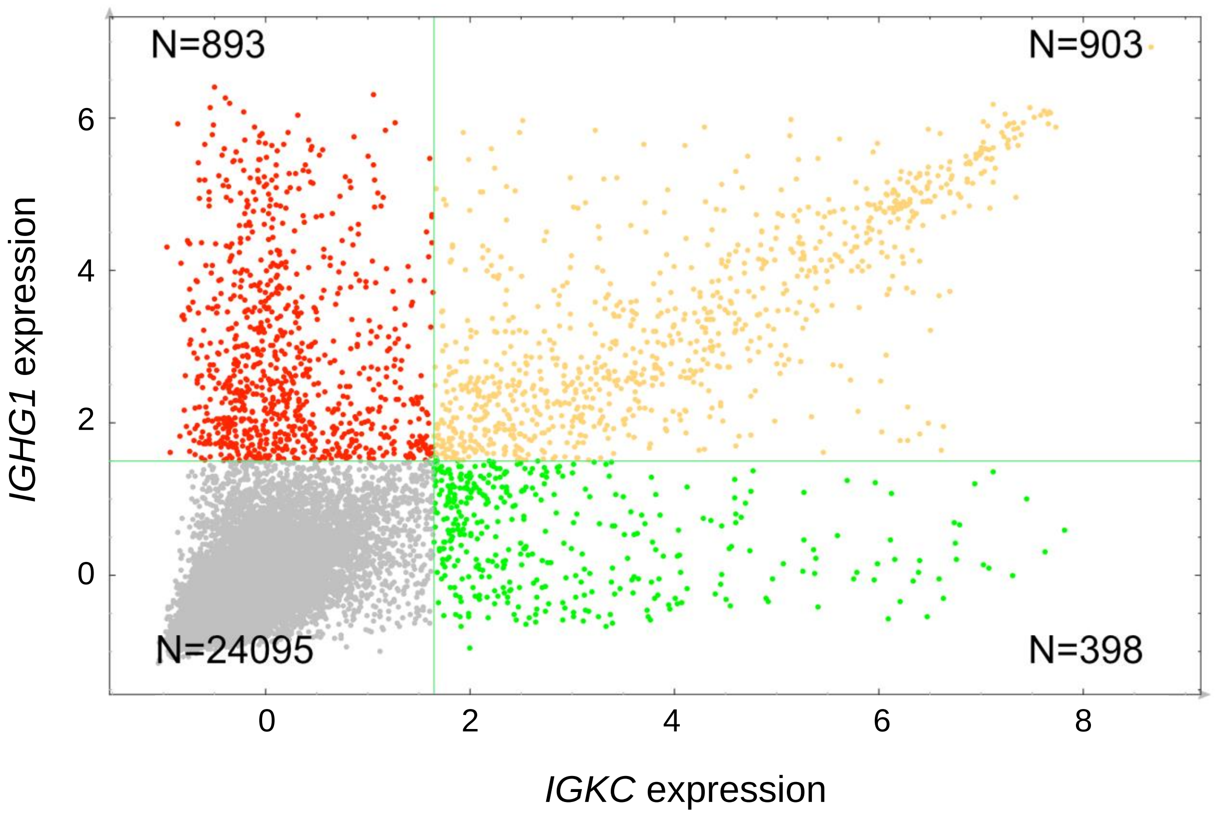


**Supplementary Fig. 7. Intensity-based quantification of *IGHG1* and *IGKC* allows for detection of plasma cells expressing different antibody heavy and light chain combinations.**
Gating of z-score normalized *IGHG1* and *IGKC* intensities.

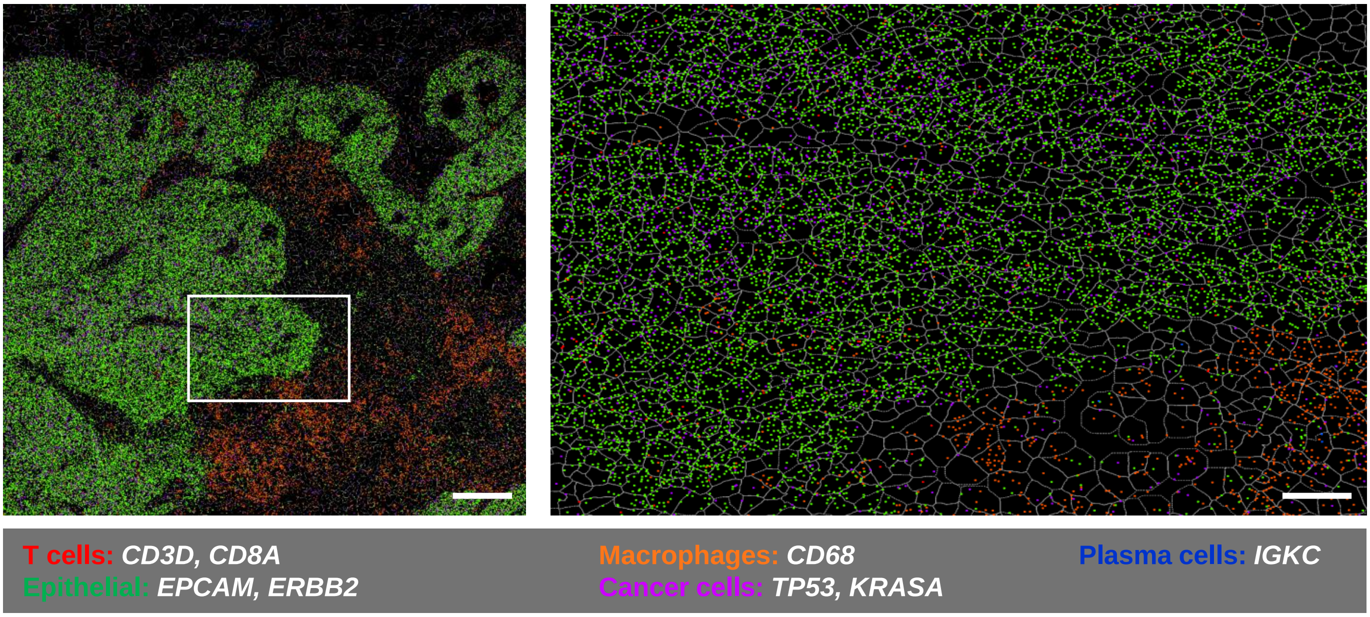


**Supplementary Fig. 8. Spatial RNA distribution in a metastatic lymph node.**
Box on left image is location of zoomed in image on right. Colored dots indicate location of select detected transcripts (legend below each image) and grey outlines show cellular segmentation. Scale bar = 200 µm; close-up region scale bar = 50 µm.

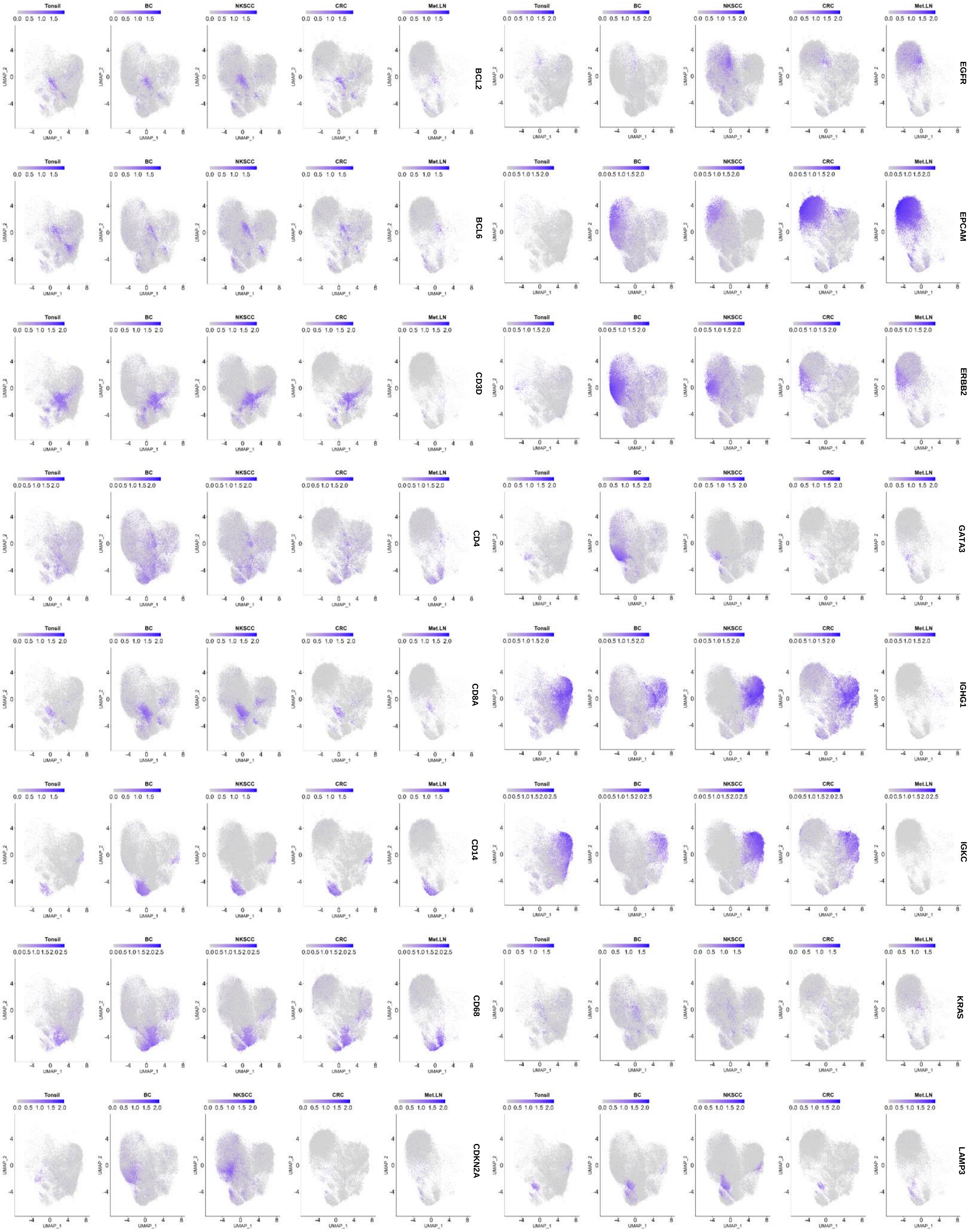
Tonsil
BC
NKSCC
CRC
Met.LN
UMAP_1
UMAP_2
BCL2
BCL6
CD3D
CD4
CD8A
CD14
CD68
CDKN2A
EGFR
EPCAM
ERBB2
GATA3
IGHG1
IGKC
KRAS
LAMP3

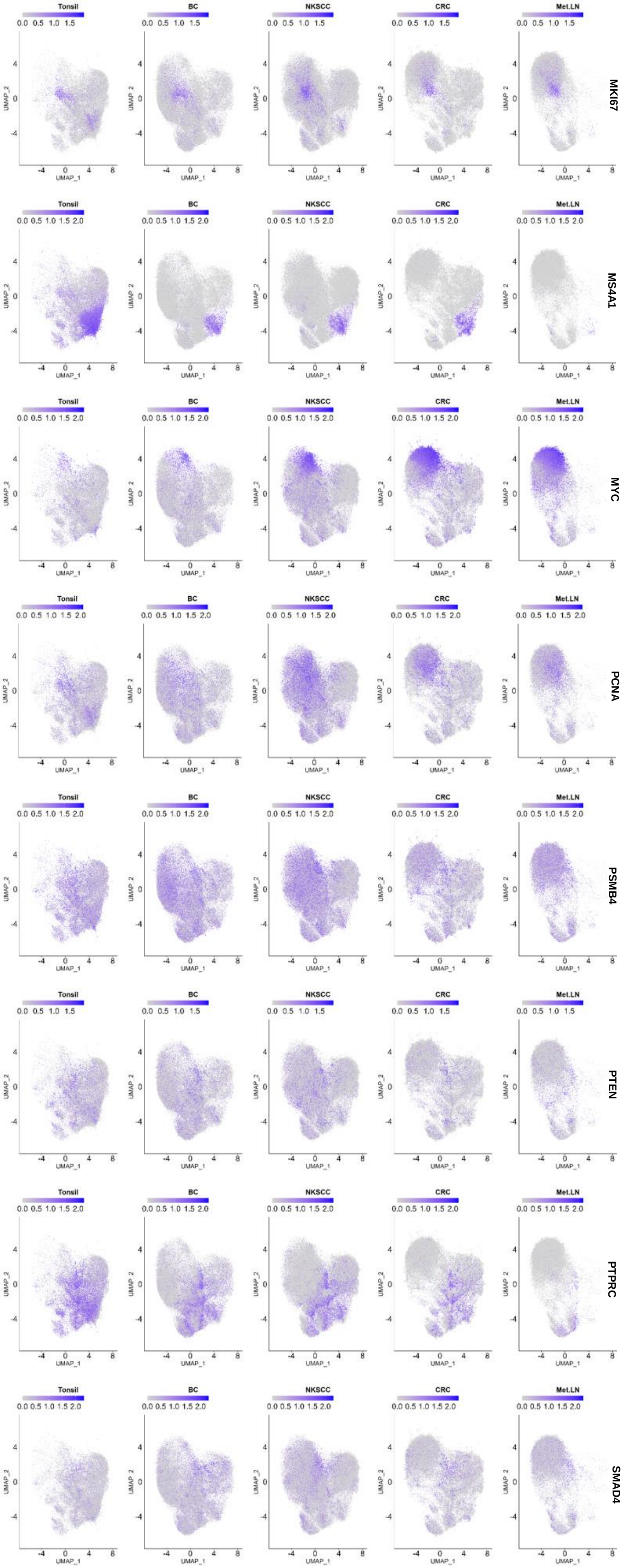


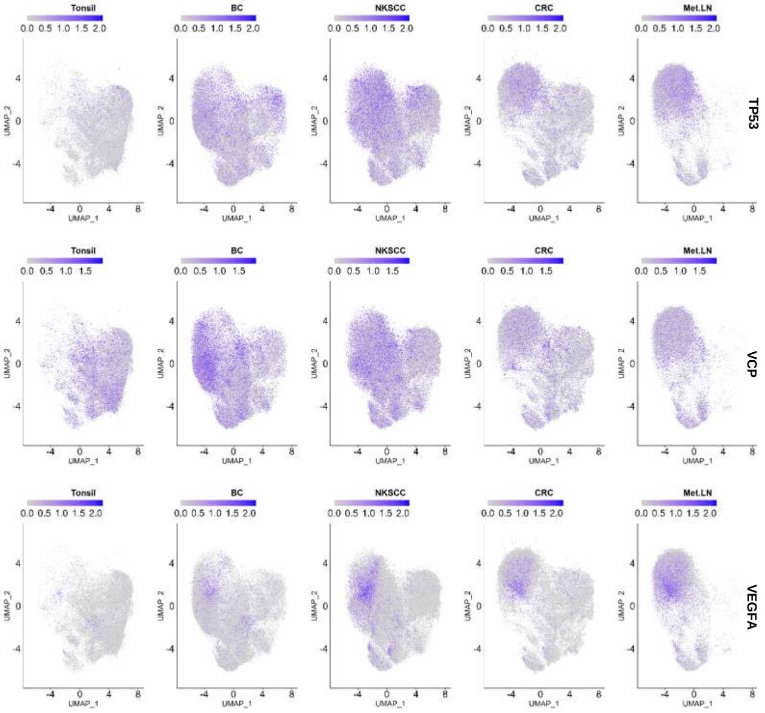


**Supplementary Fig. 9. Gene expression profiles across cancer samples.**
UMAP plots of all tissues (BC – breast cancer, NKSCC – non-keratinizing squamous cell carcinoma, CRC – colorectal cancer, Met. LN – metastatic lymph node) analyzed exhibiting expression patterns of each gene in the extended I-O panel across samples.

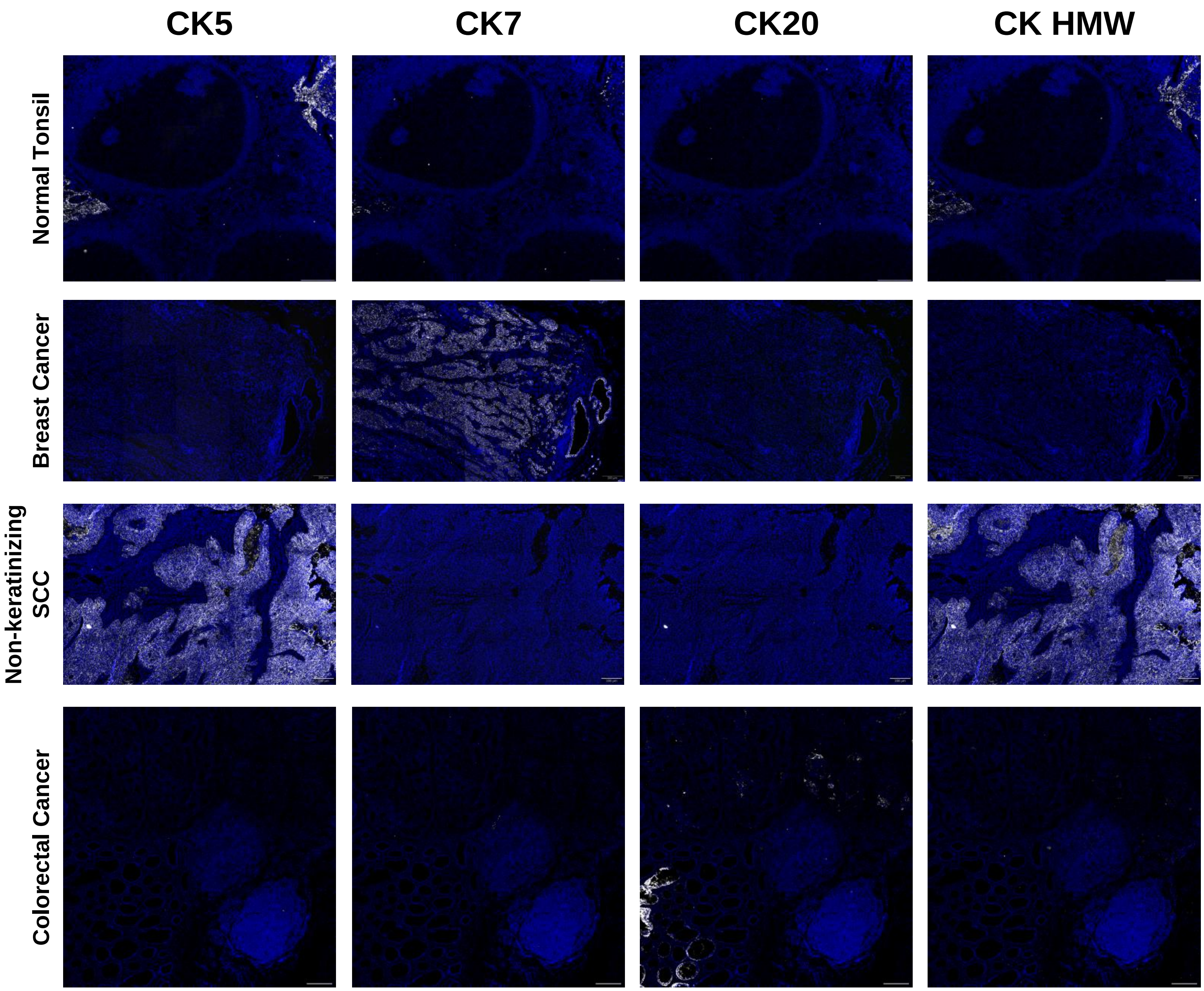


**Supplementary Fig. 10. Immunostaining of cytokeratins for pathological tissue assessment.** Immunofluorescence images of selected cytokeratins. Scale bar = 200 µm.

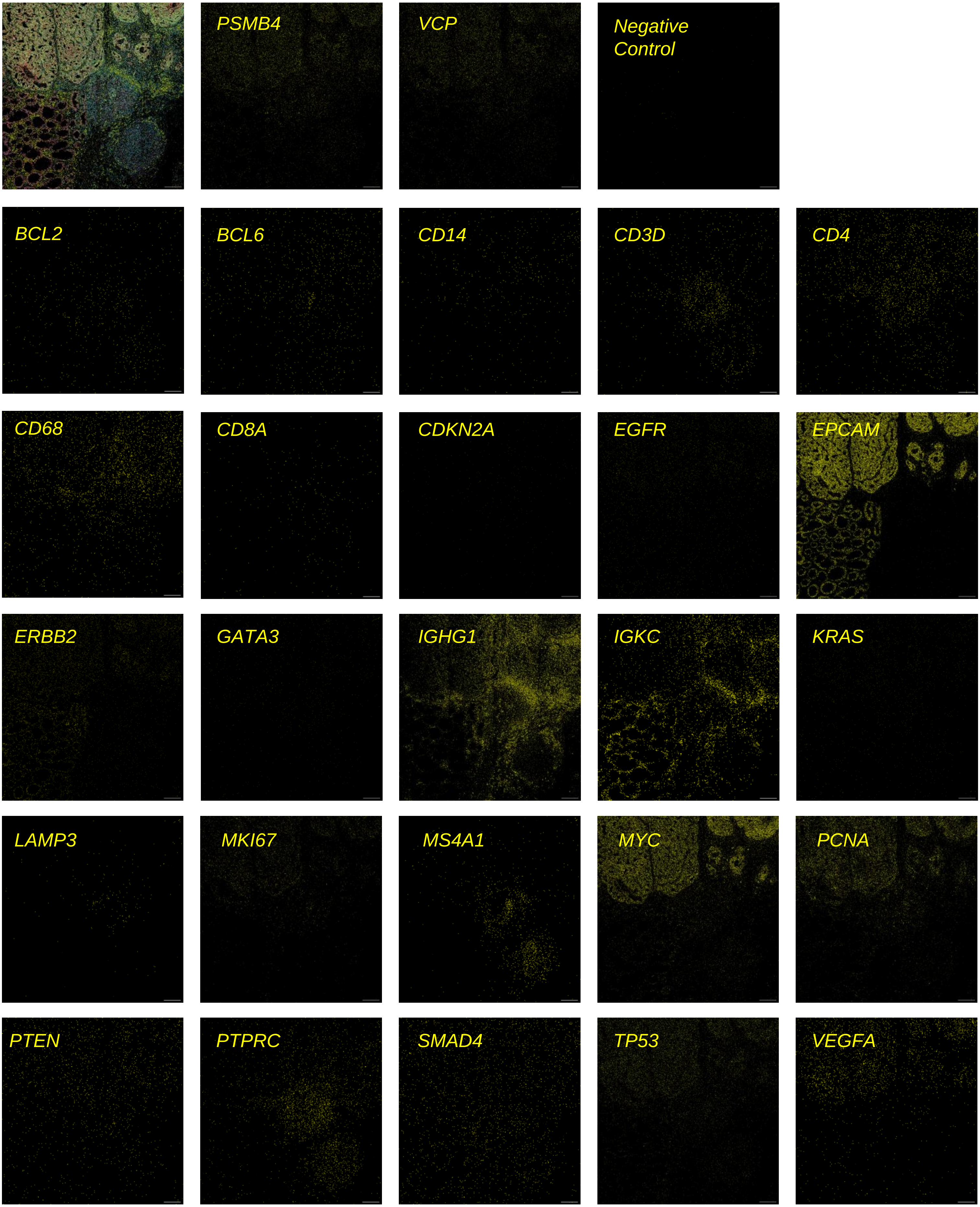


**Supplementary Fig. 11. Spatial distribution of individual genes in a colorectal cancer.** Location of RNA transcripts detected by RNAsky for each gene in the extended I-O panel set in Fig. 4c. Housekeeping genes (*PSMB4* and *VCP*), negative control, and combined image of all transcripts detected in top row. Scale bar = 200 µm.

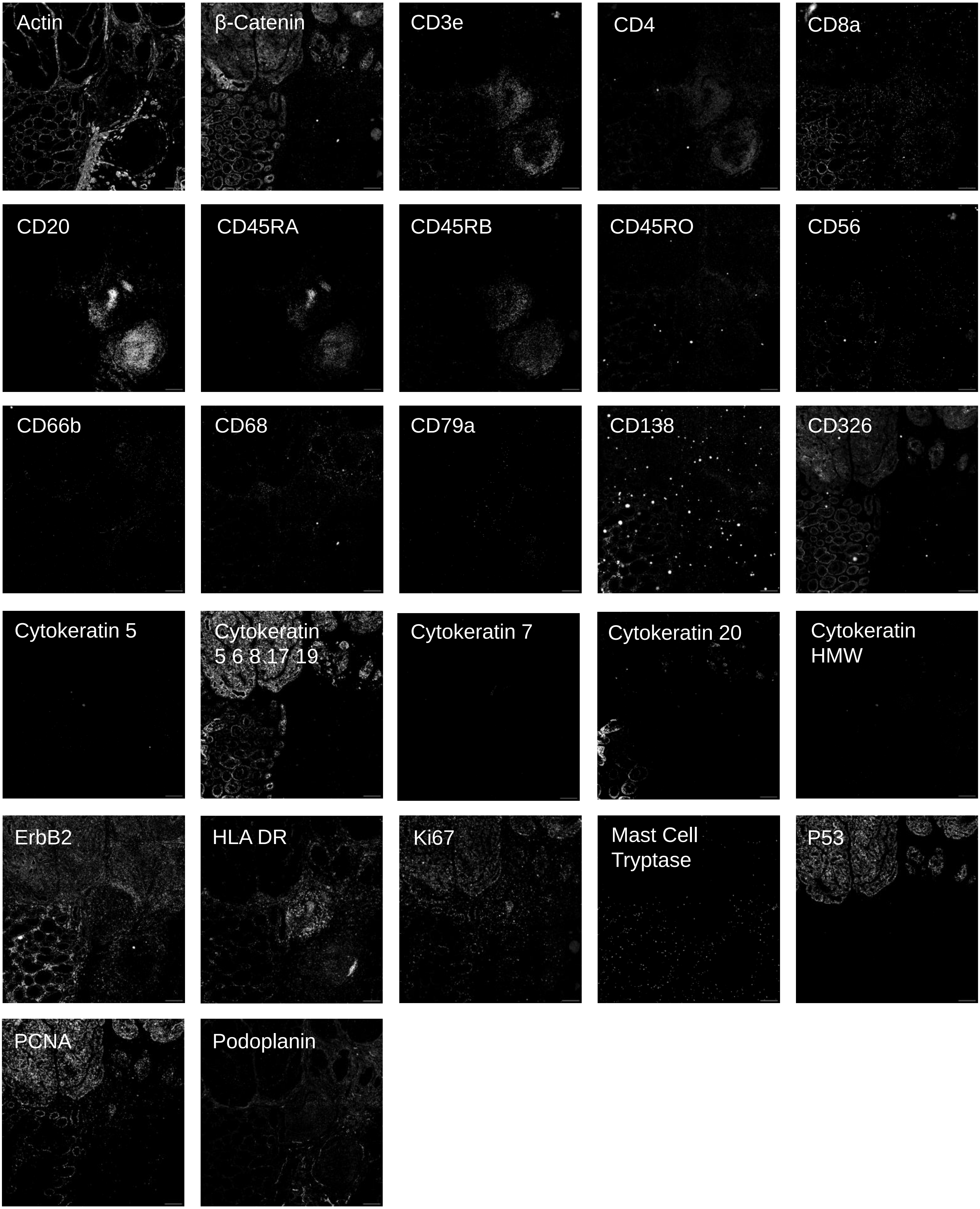


**Supplementary Fig. 12. Immunostaining of individual proteins in a colorectal cancer.** Immunofluorescence image of each protein assayed. Scale bar = 200 µm.

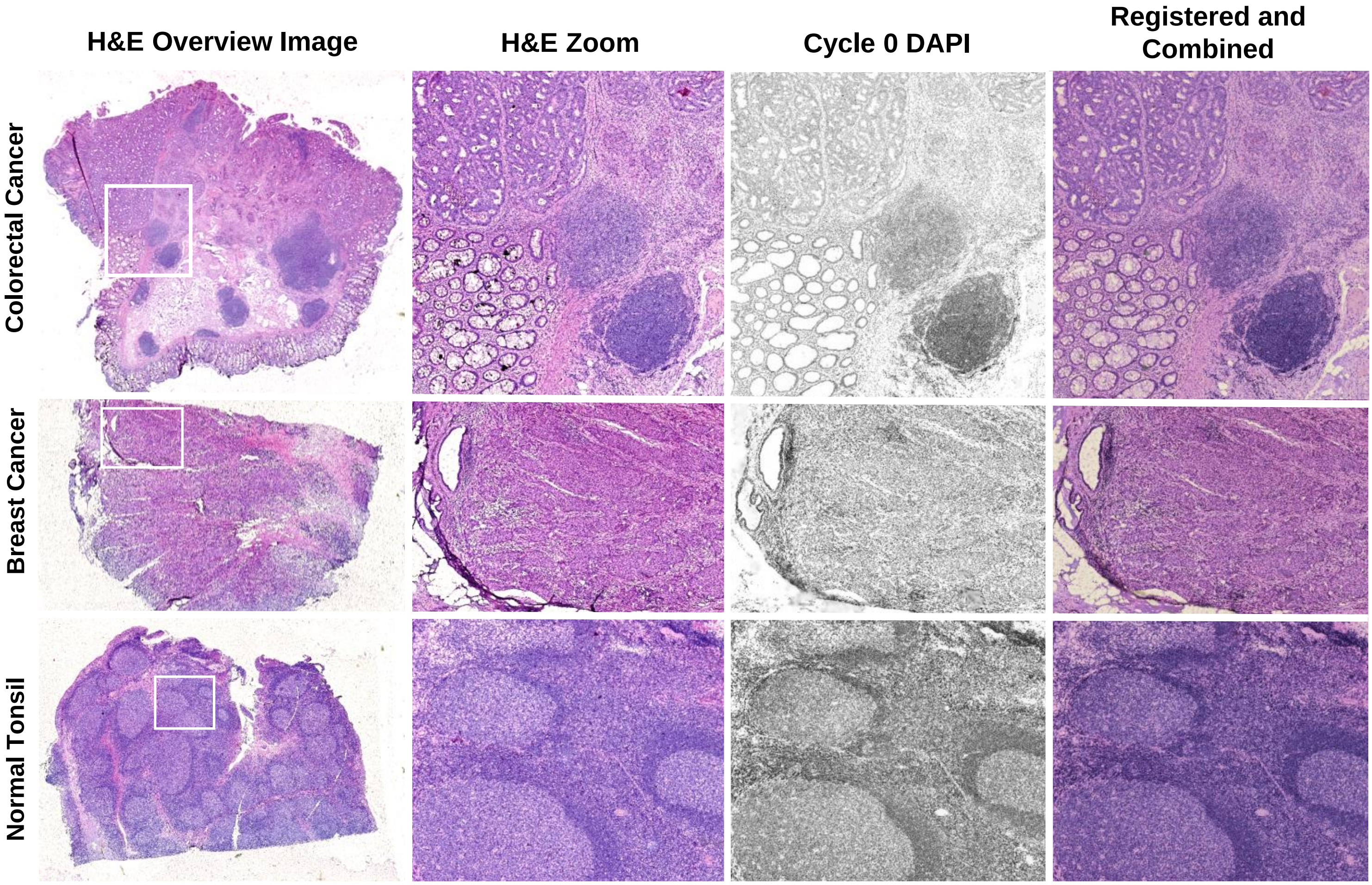


**Supplementary Fig. 13. Registration of H&E image to DAPI channel.**
Left column shows H&E image of whole tissues (CRC – colorectal cancer, BC – breast cancer) following an RNAsky and immunostaining MACSima run. White boxes indicate ROIs selected for the run and downstream analysis. Middle columns show separate H&E and dapi staining, respectively for selected ROIs. Right column shows the registered and combined H&E and dapi stained images.
